# Not4-dependent targeting of *MMF1* mRNA to mitochondria limits its expression via ribosome pausing, Egd1 ubiquitination, Caf130, No-Go-Decay and autophagy

**DOI:** 10.1101/2022.08.29.502450

**Authors:** Siyu Chen, George Allen, Olesya O. Panasenko, Martine A. Collart

**Affiliations:** Department of Microbiology and Molecular Medicine, Faculty of Medicine, University of Geneva, Institute of Genetics and Genomics of Geneva, Geneva, Switzerland

**Keywords:** Ccr4-Not complex, Not4, Egd1, Mitochondria, co-translational import, *MMF1*, Caf130, targeted No-Go-Decay, autophagy

## Abstract

The Ccr4-Not complex is a conserved multi protein complex with diverse roles in the mRNA life cycle. Recently we determined that the Not1 and Not4 subunits of Ccr4-Not inversely regulate mRNA solubility and thereby impact dynamics of co-translation events. One mRNA whose solubility is limited by Not4 is *MMF1* encoding a mitochondrial matrix protein. In this work we determine that Not4 promotes the co-translational docking of *MMF1* mRNA to mitochondria via the mitochondrial targeting sequence of the Mmf1 nascent chain, the Egd1 chaperone, the Om14 mitochondrial outer membrane protein and the co-translational import machinery. We observe that *MMF1* mRNA is translated with ribosome pausing and uncover a mechanism that depends upon its targeting to the mitochondria and limits its overexpression. We have named this mechanism Mito-ENCay. It relies on **E**gd1 ubiquitination by **N**ot4, the **C**af130 subunit of the Ccr4-Not complex, the mitochondrial outer membrane protein **C**is1, No-Go-Dec**ay** as well as autophag**y**. We propose that in fermenting yeast, mRNAs whose encoded proteins depend upon co-translational folding and/or assembly are regulated by Caf130-dependent quality control mechanisms similar to Mito-ENCay.

## Introduction

Mitochondria are essential organelles with functions in cellular metabolism and homeostasis. They are of central importance for cellular energetics and participate in signaling mechanisms that ensure survival or promote death of cells under stress (1,2). Disruption of mitochondrial function has been associated with a large variety of diseases (3,4). Mitochondria have a characteristic architecture, delimited by outer and inner membranes, with inner membrane invaginations called cristae where oxidative phosphorylation occurs. The inner most aqueous compartment is the matrix. More than 1000 proteins have been identified in yeast mitochondria and nuclear genes encode over 99% of these. Hence, mitochondrial precursor proteins are for the most part produced in the cytoplasm and must be targeted to the appropriate mitochondrial compartments by targeting signals. In some cases the mitochondrial mRNAs are targeted to the mitochondria where they are translated and proteins co-translationally imported ((5–8) and for review see (9)), while in other cases proteins are synthesized in the cytosol and must reach the mitochondria post-translationally. Little is known about how such proteins reach the mitochondria *in vivo* (10). Targeting of the mRNAs to the mitochondria can be mediated by RNA binding proteins associating with 3’ untranslated regions (UTR) independently of translation, or by the mitochondrial targeting sequence of the nascent chains during translation. In budding yeast, the Puf3 RNA binding protein has important roles in targeting mitochondrial-specific mRNAs to the surface of mitochondria in respiratory conditions (6,7,11). For translation-dependent targeting, mitochondrial mRNAs can rely on the Egd1 subunit of the NAC chaperone, the Om14 mitochondrial outer membrane (MOM) protein and Tom20 of the import machinery (8,12). The NAC chaperone is a heterodimer composed of alpha (Egd2 in yeast) and beta (Egd1 or Btt1 in yeast) subunits and it binds nascent peptides during translation (13,14). It is present in polysomes producing nuclear-encoded mitochondrial mRNAs (15,16). In all cases, the mitochondrial protein import machineries must take up the mitochondrial precursor proteins. These machineries are diverse and at least five major import pathways have been identified so far, each pathway characterized by a different machinery and different targeting signals (for review see (17)).

Important quality control (QC) systems respond to overexpressed mitochondrial precursors, to aberrant, mis-targeted or stalled nascent proteins at the MOM, to a saturated or compromised import channel, but also to excessive aggregated proteins in the cytoplasm, that all collaborate to maintain cellular homeostasis (for review see (18)). Nascent chains stalled on the ribosome and engaged with mitochondrial import channels are rescued by the ribosome-associated quality control (RQC) complex, comprised of the Ltn1 ubiquitin ligase, the ATPase Cdc48, Rqc1 and Rqc2. RQC assembles on the 60S ribosomes containing unreleased peptidyl-tRNA. Vms1, a tRNA hydrolase that releases the stalled polypeptide chains engaged by the RQC (19), antagonizes Rqc2 to prevent elongation of the nascent chain with carboxy-terminal alanyl/threonyl (CAT) tails. Thereby it facilitates the import and degradation of the nascent chains in mitochondria (20). Instead, CAT-tail extension by Rqc2 ensures ubiquitination of stalled nascent chains by Ltn1 for degradation by the proteasome in the cytosol. Aberrant accumulation of mitochondrial precursors in the cytosol leads to a stress response that has been termed “mPos” that can be attenuated by a feedback loop involving changes in specific gene expression and protein chaperoning (Wang et al., 2015). Defects in protein import is one way by which an accumulation of mitochondrial precursor proteins can occur. “Mito-TAD”, is a response in which Ubx2 clears trapped precursor proteins from the TOM channel under non-stress conditions (Martensson et al., 2019). “MitoCPR” is a response that facilitates degradation in the cytosol of unimported mitochondrial precursor proteins accumulating at the mitochondrial translocase. It involves inducing expression of Cis1 at the translocase, that functions with the AAA^+^ adenosine triphosphatase Msp1 and the proteasome (Weinberg and Amon, 2018). This improves mitochondrial import during import stress. Under these conditions, some mitochondrial proteins also get degraded in the nucleus by “mitoNUC”. This process is mediated by the combined action of the E3 ubiquitin ligases San1, Ubr1 and Doa10 and requires an N-terminal mitochondrial targeting sequence (Shakya et al., 2021). “MAD” is the response by which the components of the ubiquitin proteasome system (UPS) are recruited to the MOM to trigger degradation of proteins peripherally associated with the MOM, integral MOM proteins, mitochondrial intermembrane space proteins, and potentially also inner membrane or matrix proteins (Braun and Westermann, 2017). An increase of mitochondrial precursor proteins in the cytosol triggers the “UPR^am^”, leading to increased proteasome assembly by the enhanced activity of the proteasome assembly factors Irc25 and Poc4, that degrades excess proteins (Wrobel et al., 2015). Inversely, upon accumulation of high levels of aggregated proteins in the cytoplasm, Hsp104 helps to dissociate the aggregates. Thereby it contributes to “MAGIC”, a mechanism by which aggregation-prone proteins can enter via import channels the mitochondrial intermembrane space or matrix for degradation (Ruan et al., 2017). All of these mechanisms are indicative of a major cross-talk between the cytoplasm and the mitochondrion to maintain protein homeostasis. In addition to these mechanisms, autophagy can sequester and remove unnecessary or dysfunctional components in bulk from the cytoplasm and mitophagy is the specific form of autophagy that serves to remove damaged mitochondria (for review see (21)).

Ccr4-Not is a conserved, multi-protein subunit complex that plays multiple roles in the control of gene expression and mRNA metabolism. In yeast Ccr4-Not consists of 9 subunits: Ccr4, Caf1, Caf40, Caf130, and the five Not proteins (Not1, Not2, Not3, Not4 and Not5) (22–25). Our current knowledge about the functional roles of this complex is that its regulatory functions span the entire lifespan of mRNAs, from their synthesis to their decay. Moreover, it plays extensive roles in translation and protein turnover (26–28). Recent studies have uncovered key roles of the Not proteins in co-translational processes, such as co-translational assembly of proteins (27,29,30) and translation elongation dynamics (31). Not5 can associate with the E site of posttranslocation ribosomes bearing an empty A site. This has been proposed to enable the Ccr4-Not complex to monitor the translating ribosome for mRNA turnover according to codon optimality (32). Consistently, depletion of Not5 changes A-site ribosome dwelling occupancies inversely to codon optimality (33). In addition, ubiquitination of Rps7A by Not4 can contribute to degradation of mRNAs by No-Go-Decay (NGD) in conditions where the RQC response is defective (34).

Recently, we noted that Not1 and Not4 depletions inversely modulated mRNA solubility thereby determining dynamics of co-translation events (https://doi.org/10.1101/2022.03.14.484207). Notably, mRNAs encoding mitochondrial proteins were enriched amongst mRNAs whose solubility was most extremely inversely regulated upon Not1 and Not4 depletion. In this context, it is interesting to note that the Ccr4-Not complex interacts with factors that contribute to targeting of mitochondrial mRNAs to the mitochondria: Egd1 is ubiquitinated by Not4 (35) and Puf3 recruits the Ccr4-Not complex to its target mRNAs for degradation (36–39). Moreover, mitochondrial mRNAs are enriched amongst mRNAs bound by Not1 in a Not5-dependent manner (28).

In our current study we uncover an integrated QC mechanism that limits levels of a mitochondrial mRNA co-translationally and mobilizes components of several of the QC systems linking cytoplasm and mitochondria described above as well as Ccr4-Not subunits. We focused our attention on one nuclear-encoded mitochondrial mRNA, *MMF1*, more soluble upon Not4 depletion (https://doi.org/10.1101/2022.03.14.484207). *MMF1* encodes a mitochondrial matrix protein required for transamination of isoleucine and it couples amino acid metabolism to mitochondrial DNA maintenance (Ernst and Downs, 2018). It forms a homotrimer proposed to interact with a trimer of Mam33 (40), a translational activator in yeast mitochondria (41). We determine that Not4 limits Mmf1 overexpression during fermentative growth by promoting the co-translational docking of its mRNA to mitochondria via the mitochondrial targeting sequence of the Mmf1 nascent chain, Egd1 and the co-translational import machinery. Accumulation of excessive *MMF1* mRNA, Mmf1 precursor and mature Mmf1 protein is then avoided in a mechanism requiring Egd1 ubiquitination by Not4, Caf130, RQC and NGD, Hsp104, as well as autophagy, a mechanism that we have called Mito-ENCay. We additionally note that in fermenting yeast the physiological targets of the Caf130 QC pathway may mostly not be mRNAs encoding mitochondrial proteins, but instead highly expressed mRNAs whose protein products are critically dependent upon co-translational protein folding and interactions to prevent their aggregation.

## Materials and Methods

### Yeast strains and plasmids

The strains, oligos, plasmids and antibodies used in this study are listed in **Table S4**. Yeast strains were grown in rich medium with 2% glucose (YPD) or in synthetic drop out medium selective for plasmid maintenance. For copper induction, cells were grown to exponential phase after dilution of an overnight culture to OD_600_ of 0.3 and a stock solution of 0.1 M CuS04 was added to a final concentration of 0.1 mM. To arrest protein synthesis a stock solution of cycloheximide (CHX) was added to a final concentration of 0.1 mg/ml in the growth medium.

The plasmid pMAC1377 expressing MCP fused to mScarlet was constructed from plasmid pMAC1105 (29) digested by XhoI and co-transformed in yeast with a PCR product amplified from pE697 with oligos 1115 and 1116, followed by plasmid rescue. The reporter plasmid expressing Mmf1 fused to Flag (pMAC1211) was constructed by cloning a PCR fragment amplified with oligos 935 and 936 and genomic DNA, digested by MfeI and Not1 in pE617 digested by EcoRI and Not1. The reporter plasmid expressing Mmf1 without the MTS (pMAC1327) was made similarly, with oligos 1009 and 936, and the one with the Cox4 MTS (pMAC1328) with oligos 1010 and 936. The one expressing Cox4 (pMAC1200) was made similarly with oligos 691 and 692, except that the PCR fragment was digested by EcoRI and Not1. For both the pMAC1211 and pMAC1327 plasmids, the *URA3* marker was swapped to the *LEU2* marker by transforming pUL9 (pE24) digested with StuI and selection of Leu+ Ura- colonies, followed by plasmid rescue leading to plasmids pMAC1341 and pMAC1342. MS2 loops were added in the pMAC1211 and pMAC1327 plasmids by co-transforming into yeast the pMAC1211 and pMAC1327 plasmids digested with SacI and a PCR fragment obtained with oligos 1087 and 1088 and pE659, leading to pMAC1365 and pMAC1367. For both plasmids the *URA3* marker was swapped to the *LEU2* marker by transforming pUL9 (pE24) digested with StuI and selection of Leu+ Ura-colonies, followed by plasmid rescue leading to plasmids pMAC1390 and pMAC1391. The cells expressing Atp5-GFP were transformed with a PCR fragment amplified with DNA from strain MY6993 with oligos Not4-5’ and Not4-V4 for deletion of the *NOT4* gene by homologous recombination according to standard procedures. The strain expressing Not4-GFP from its endogenous locus (MY14341) was made with F2 and R1 oligos and pE85 by homologous recombination according to standard procedures (42). All plasmids were verified by sequencing. Plasmids encoding Egd1, Not4 and Rli1 derivatives have already been published (see **Table S4**).

### Protein ubiquitination assay

A plasmid expressing 6His-tagged ubiquitin under the control of the inducible *CUP1* promoter was transformed into cells. The transformants were cultured in medium selective for plasmid maintenance in the presence of 0.1 mM CuSO4. 100 OD_600_ of cells were harvested when they reached late exponential phase. Cell pellets were weighed and resuspended with G-buffer (100 mM sodium Pi, pH 8.0, 10 mM Tris-HCl, 6 M guanidium chloride, 5 mM imidazole, 0.1% Triton X-100) at 100 mg/ml. 0.6 ml of glass beads was added and cells were disrupted by bead beating for 15 minutes at room temperature (RT). Following centrifugation, 20 μl of the supernatant was taken as total extract (TE), and 700 μl of the supernatant was mixed with 30 μl of nickel-nitrilotriacetic acid-agarose (Ni-NTA, Qiagen) for 2 h at RT with mild rotation. U-buffer (100 mM sodium Pi, pH 6.8, 10 mM Tris-HCl, 8 M urea, 0.1% Triton X-100) was used to abundantly wash the Ni-NTA-agarose to which ubiquitinated proteins were bound. SB was added directly to the Ni-NTA with the ubiquitinated proteins for analysis by western bloting with relevant antibodies.

### Confocal Microscopy

For imaging cells were grown in selective synthetic media for plasmid selection as indicated to an OD_600_ between 0.6 and 1.2. 2 OD_600_ of cells was collected by the centrifugation at 3000 g for 5 min at RT. The cell pellets were washed twice with 1 ml PBS. The final cell pellets from 0.5 OD_600_ of cells were resuspended in 200 μl PBS, 20 μl of which was loaded on 1 % agarose gel coated coverslips, with an even distribution of cells. Then the coverslips were mounted by nail polish. The prepared slides were immediately imaged with a standard confocal microscope (LSM800 Airyscan) with a 63 X oil objective (NA=1.4) that was used for image acquisition. Each image was acquired by z-stacking. The image J software was used to process the images for colocalization analysis and for this co-localization analysis, more than 20 cells of each sample were evaluated. The acquired Pearson’s R value of co-localization was statistically analyzed by Prism9.

### Protein extracts, SDS-or Native PAGE and Western blotting

Total protein extracts were prepared by incubating pelleted yeast cells in 0.1 M NaOH for 10 min at RT. After a quick spin in a microfuge, the cell pellet was resuspended in 2 X sample buffer (post-alkaline lysis). Samples were subjected to SDS-PAGE and western blotting according to standard procedures. For native gels, ready-made native 3–12% Bis-Tris gels were used (Invitrogen) according to instructions. Briefly, 20 OD_600_ of cells were harvested at exponential growth. Cells were disrupted by 0.2 ml glass beads in the presence of 0.4 ml lysis buffer (20 mM Hepes pH 7.5, 20 mM KCl, 10 mM MgCl2, 1 % Triton X-100, 1 mM DTT, 1 mM PMSF, supplemented with a cocktail of protease inhibitors (Roche)). The indicated amount of total protein extract was mixed with native sample buffer from Invitrogen. Following the electrophoresis (150 V, 3 h, 4 °C) and transfer (40 W, 1 h, RT) to PVDF membranes, the blots were incubated with the indicated antibodies (43).

### Sedimentation through a sucrose cushion

For polysome sedimentation 100 OD_600_ of cells were harvested at exponential growth. Cells were disrupted by 0.2 ml glass beads in the presence of 0.4 ml lysis buffer (20 mM Hepes pH 7.5, 20 mM KCl, 10 mM MgCl2, 1 % Triton X-100, 1 mM DTT, 1 mM PMSF, supplemented with a cocktail of protease inhibitors). 20 μl of lysate was taken as input control; 200 μl of the remaining lysate was loaded onto a 60 % sucrose cushion in 0.5 ml mini-ultracentrifuge tubes. Following ultracentrifugation (85000 rpm, 90 min, 4°C) in a Sorvall S120-AT2 Fixed-Angle Micro-Ultraspeed Rotor, the pellet at the bottom of the mini-ultracentrifuge tube was resuspended with 200 μl of lysis buffer. The resuspended pellet was analyzed by SDS-PAGE and western blotting.

### Tandem affinity purification and mass spectrometry

Not1-Taptag, Not5-Taptag, and Caf1-Taptag were purified by tandem affinity purification from wild type and *caf130Δ* cells as previously described (44). The purified proteins were identified by LC-MS/MS at the proteomics platform of the Faculty of Medicine (https://www.unige.ch/medecine/proteomique/). ESI LC-MS/MS was performed on LTQ Orbitrap velos (Thermo Fisher Scientific) equipped with a NanoAcquity (Waters). Peptides were trapped on a home-made 5 μm 200 Å Magic C18 AQ (Michrom) 0.1 × 20 mm pre-column and separated on a home-made 5 μm 100 Å Magic C18 AQ (Michrom) 0.75 × 150 mm column with a gravity-pulled emitter. The analytical separation was run for 35 min using a gradient of H2O/FA 99.9%/0.1% (solvent A) and CH3CN/FA 99.9%/0.1% (solvent B). The gradient was run as follows: from 0 to 65% A in 14 min., and 20% then to 80% B in 5 min at a flow rate of 220 nL/min. For MS survey scans, the OT resolution was set to 60000 and the ion population was set to 5 × 105 with an m/z range from 400 to 2000. Five precursor ions were selected for collision-induced dissociation (CID) in the LTQ. For this, the ion population was set to 7×103 (isolation width of 2 m/z). The normalized collision energies were set to 35% for CID. Peaklists (MGF file format) were generated from raw data using the MS Convert conversion tool from ProteoWizard. The peaklist files were searched against the *Saccharomyces cerevisiae* database (UniProtKB) and with an in-house database of common contaminants using Mascot (Matrix Science, London, UK). Trypsin was selected as the enzyme, with one potential missed cleavage. Precursor ion tolerance was set to 10 ppm and fragment ion tolerance to 0.6 Da. Variable amino acid modifications were oxidized methionine, fixed amino acid modification was carbamidomethyl cysteine. The Mascot search was validated using Scaffold (Proteome Software). Peptide identifications were accepted if they could be established at greater than 95.0% probability by the Peptide Prophet algorithm (45). Protein identifications were accepted if they could be established at greater than 95.0% probability and contained at least 2 identified peptides. Protein probabilities were assigned by the Protein Prophet algorithm (46). Proteins that contained similar peptides and could not be differentiated based on MS/MS analysis alone were grouped to satisfy the principles of parsimony. Summed NSAF values for each sample were kept for further analysis.

### RNA preparation and analysis

RNA extraction and analysis was performed as previously described (28). Relative mRNA abundances were determined by RT-qPCR with the Pfaffl method (47). For normalization, we measured *EGD2* as an invariable control mRNA and calculated the ΔCT values.

### Mitochondria isolation

The Mitochondrial Yeast Isolation kit and protocol (ab178779; Abcam) were used to fractionate yeast cells by differential centrifugation. Briefly, cells with the different reporters were grown in synthetic drop-out media at 30 °C until the logarithmic growth phase. 25 OD_600_ of cells were collected by centrifugation (3000 g, 5 min, RT). Cell pellets were resuspended with the buffer A containing 10 mM DTT, then buffer B containing Lysis Enzyme Mix provided by the supplier, to form spheroplasts. From this step onwards, the procedure was on ice. The spheroplasted cells were resuspended by homogenization in Buffer C provided by the supplier + a protease inhibitor cocktail provided, transferred to a glass douncer, and broken by 10 –15 strokes. 20 μl of lysate was put aside as input. Following centrifugation at 600 g for 5 min at 4°C, the supernatant, which contained the intact mitochondria, was collected. Further centrifugation of the supernatant (12000 g, 10 min, 4 °C), led to a sedimented fraction containing mitochondria. 600 μl of the supernatant at this step (cytoplasm) was taken and mixed with 120 μl 100% trichloroacetic acid solution and incubated for 10 min at 4°C to precipitate the cytoplasmic proteins. The input, mitochondrial and cytoplasmic pellet fractions were mixed with 2 X SB and analyzed by western blotting.

### Ribosome profiling and bioinformatic analysis

Samples for ribosome profiling were prepared as described previously (30). For the analysis of the Ribo-Seq samples, all fastq files were adaptor stripped using cutadapt (48). Only trimmed reads were retained, with a minimum length of 20 and a quality cutoff of 2 (parameters: -a 10 CTGTAGGCACCATCAATAGATCGGAAGAGCACACGTCTGAACTCCAGTCAC–trimmed-only–minimum-length = 20–quality-cutoff = 2). Histograms were produced of ribosome footprint (RPF) lengths that were very homogeneous with highest reads between 28 and 31 that were kept for the analysis. Reads were mapped, using default parameters, with HISAT2 (49) to R64-1-1, using Ensembl release 84 gtf for transcript definitions. UTR definitions were taken from the *Saccharomyces* Genome Database and a standard region of 100bp was used where a gene’s UTR was not defined. A minimum length of 30bp was implemented to ensure appropriate mapping around the start and stop codons. For the mapping, only unique alignments to transcripts were retained. A full set of 6692 CDSs were established for R64-1-1 Ensembl release 84 and extended by the same UTR sequences defined above. The filtered reads were then mapped to this transcriptome with bowtie2 (50), using default parameters. For all downstream analysis, dubious ORFs were filtered to leave 5929 transcripts. The A/P site position of each read was predicted by riboWaltz (51) and aggregated over all transcripts. Differential expression was performed using DESeq2 on default settings (52) and enrichment tests were performed using the ‘phyper’ hypergeometric test in R with GO Slim gene set definitions. A CDS was considered to have a large pause in *caf130*Δ if the P-site depth (per million genome-wide) in one codon position was 3 or above, 2-fold more than the same position in WT and greater than 5 standard deviations higher than the mean normalised depth over the whole CDS (excluding codons with no coverage) in *caf130*Δ. The new data is accessible online as GSE206973.

### RNA-Seq and Solubility analyses

The data was generated and analyzed in (33). To briefly describe its origin, total and soluble RNA were isolated from cells with Not1 and Not4 degron alleles before and after auxin treatment to deplete Not1 and Not4. The same amount of RNA from each sample was spiked in with a same amount of *S.Pombe* RNA, libraries were generated and sequenced. Sequencing files were demultiplexed using bcl2fastq v2.20.0.422 (one mismatch, minimum length 35 nt), and adapters were trimmed using cutadapt 2.3. (48) at default settings, allowing one mismatch and minimum read length of 35nt. In addition to standard Illumina dual index (i5, i7), the inline sample and UMI barcode was analyzed using Umitools. Reads were mapped to the concatenated genome of *S. cerevisiae* (R64-1-1) and *S. pombe* (ASM294v2) using STAR. CDS positions were defined with Ensembl gff version 94 for of *S. cerevisiae* (R64-1-1). Counts in *S. cerevisiae* were calculated by aggregating RNA-Seq reads overlapping CDS positions. Differential expression was performed using DESeq2 on default settings (52). We then define solubility as the log fold change produced by DESeq2, comparing RNA-Seq counts for the soluble fraction in a given sample by the corresponding counts for the total fraction of the same sample as described previously (33).

## Results

### Mmf1, but not Cox4, is co-translationally imported

To start dissecting how the Ccr4-Not complex regulates solubility of mRNAs to regulate co-translation events, we focused our attention on two mitochondrial mRNAs, *MMF1* and *COX4*, rendered more soluble upon Not4 depletion, but less soluble upon Not1 depletion (**Figure S1A**). Both mRNAs express mitochondrial precursor proteins with an N-terminal cleavable targeting sequence and assemble into multi-protein complexes. However, Cox4 is a component of the respiratory complex IV located in the mitochondrial inner membrane whereas Mmf1 resides in the matrix. Moreover, the *COX4* and *MMF1* mRNAs differ by their ribosome profiling data in wild type cells (30) showing probable ribosome pausing for *MMF1* but not for *COX4*, though ribosome footprints are increased for both *COX4* and *MMF1* mRNAs in *not4Δ* (31) (**Figure S1B**). Interestingly, for *MMF1* the increase is mostly after the pause site (**Figure S1C**), suggesting decreased efficiency of ribosome pausing in *not4Δ*.

To study the regulation of *MMF1* and *COX4* expression dependent upon their coding sequences, and the role of Not4, we used reporter constructs with the heterologous and inducible *CUP1* promoter and the heterologous *ADH1* 3’UTR in between which we cloned the *MMF1* and *COX4* coding sequences (CDS) fused to a C-terminal Flag tag (**Figure 1A**). We transformed the plasmids in wild type cells and tested expression of the reporter before and after induction with copper for 10 min. Before induction some mature Mmf1 was already detectable, due to some leakage of the *CUP1* promoter. Immediately after induction, levels of unprocessed and mostly mature Mmf1 were increased, whilst mostly unprocessed Cox4 was visible, with very low levels of mature protein (**Figure 1B**). This suggests that processing of induced Mmf1 might be faster than that of Cox4, compatible with the idea that the former but not the latter might be co-translationally processed and imported.

**Figure 1.**
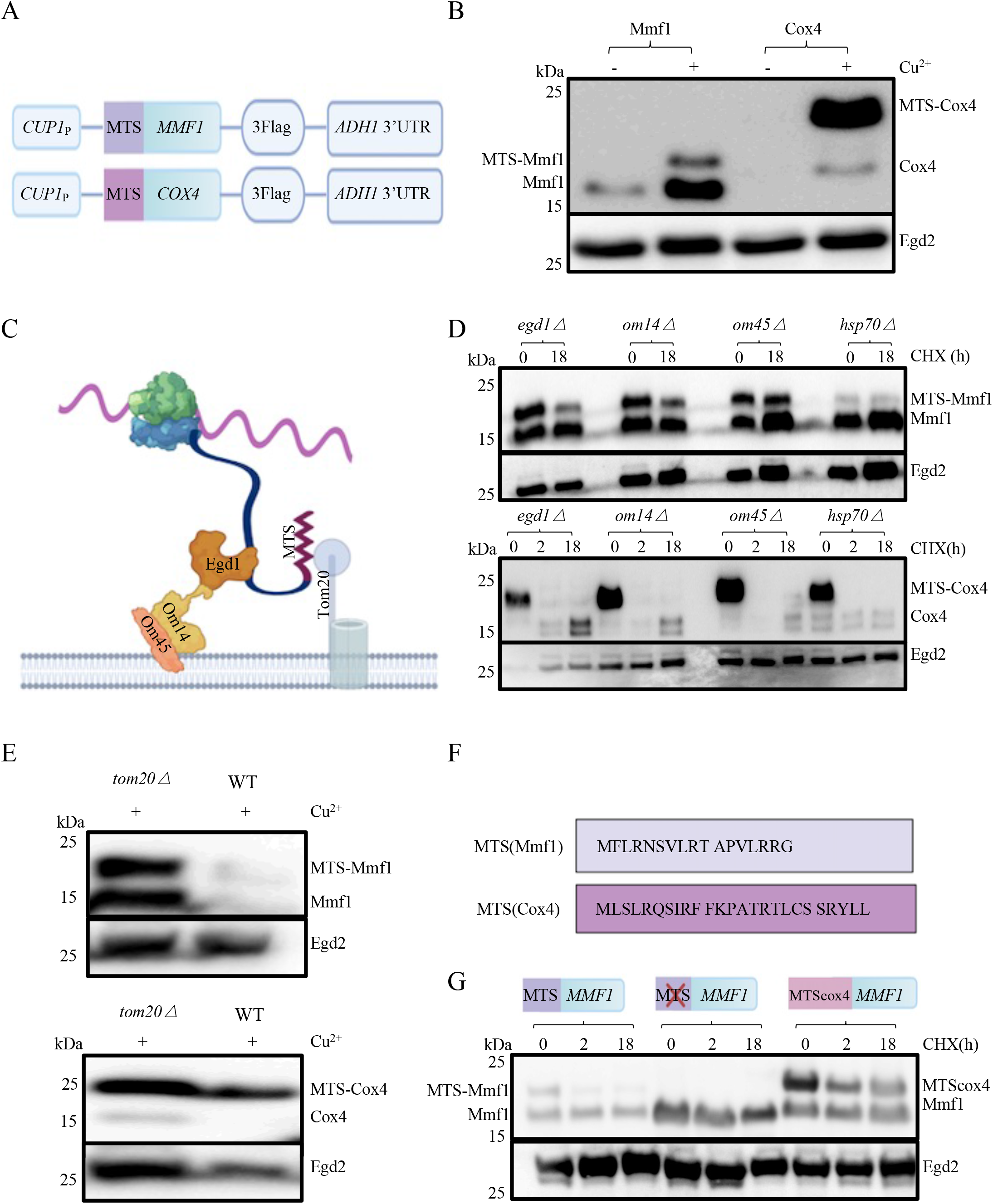
Mmf1 but not Cox4 is co-translationally imported and regulated. **A**. Cartoon of the reporter constructs used in which coding sequences are fused to a C-terminal Flag tag, under the control of the *CUP1* inducible promoter. **B**. Wild type cells (WT) transformed with the reporters and growing exponentially in medium selective for the plasmids were untreated (-) or treated (+) with 0.1 mM CuS04 (Cu^2+^) for 10 min. Cells were collected for total protein analysis by western blotting with antibodies to Flag or with antibodies to Egd2 to control for protein loading. Precursor and mature Mmf1 and Cox4 are indicated respectively left and right of the blot. Molecular weight markers are indicated on the left. **C**. Cartoon of the co-translational import machinery with the nascent chain exposed from the ribosome interacting with the Egd1 chaperone itself docking onto the Om14 OM protein interacting with Om45, and the MTS of the nascent chain recognizing Tom20 to enable transfer of the nascent chain into the Tom channel. **D** and **E**. Analysis of the reporters as in **panel B** in the indicated mutant strains in cells after induction (0) then treated or not with CHX at 100 μg/ml and left at 30°C for 2 or 18h (as indicated). **F**. Amino acid sequence of the Mmf1 and Cox4 MTS. **G**. Analysis as in **panel B** of the *MMF1* reporter with MTS (left), without MTS (middle), or with the Cox4 MTS to replace its own MTS (right), in wild type cells.

To look at this further we transformed the 2 plasmids in strains defective for the mitochondrial co-translational import machinery, namely cells lacking the Egd1 chaperone or its receptor on the MOM, Om14, or the Om14 partner Om45, or finally the Tom20 receptor (see cartoon on **Figure 1C**). We also transformed the plasmids in cells defective for the cytoplasmic Hsp70 chaperones reported to contribute to effective post-translational import of mitochondrial proteins (53). As before, we induced the expression from the reporter plasmids for 10 min with copper, then we did a cycloheximide (CHX) chase up to 18 h to follow turnover of the induced proteins. After induction, we noted elevated levels of the unprocessed Mmf1 protein in the mutants of the co-translational machinery but not in the *hsp70* mutant (compare **Figure 1D**, upper panel, and **Figure 1B**), and the unprocessed Mmf1 mostly did not turn over in the 18 h of chase. In all strains, immediately after induction we noted mostly unprocessed Cox4 that was effectively turned over already after 2 h, such that at 18 h only low levels of mature Cox4 were detectable (**Figure 1D**, lower panel). Similarly, Mmf1 but not Cox4 was increased in the *tom20* mutant compared to the wild type strain after a 10 min copper induction (**Figure 1E**). These results are compatible with a role of the co-translational import pathway for control of Mmf1, but not Cox4, expression. Moreover, they indicate that the Mmf1 precursor is not rapidly turned over while the Cox4 precursor is.

To analyze this further, we investigated the role of the mitochondrial targeting sequence (MTS) for regulation of Mmf1 expression and the ability of the Cox4 MTS to replace the Mmf1 MTS. Indeed, both Mmf1 and Cox4 have an N-terminal cleavable MTS, but the amino acid sequence of the MTS is very different (**Figure 1F**). Mmf1 expressed without its MTS or with the Cox4 MTS to replace its own MTS was overexpressed (**Figure 1G**). These results indicate that the Mmf1 MTS is necessary to limit Mmf1 expression, and that the Cox4 and Mmf1 MTS are not functionally interchangeable for this function.

### Regulation of *MMF1* but not *COX4* expression requires Not4 and the MTS

We next tested expression and turnover of Mmf1 and Cox4 expressed from the *MMF1* and *COX4* reporters in cells lacking Not4. The expression of the Mmf1 precursor and mature protein was much higher in *not4Δ* right after induction, and both the processed and unprocessed Mmf1 stayed high after the CHX chase (**Figure 2A**, upper panels). In contrast the expression of Cox4 was mostly indistinguishable between the wild type and mutant (**Figure 2A**, lower panels).

**Figure 2.**
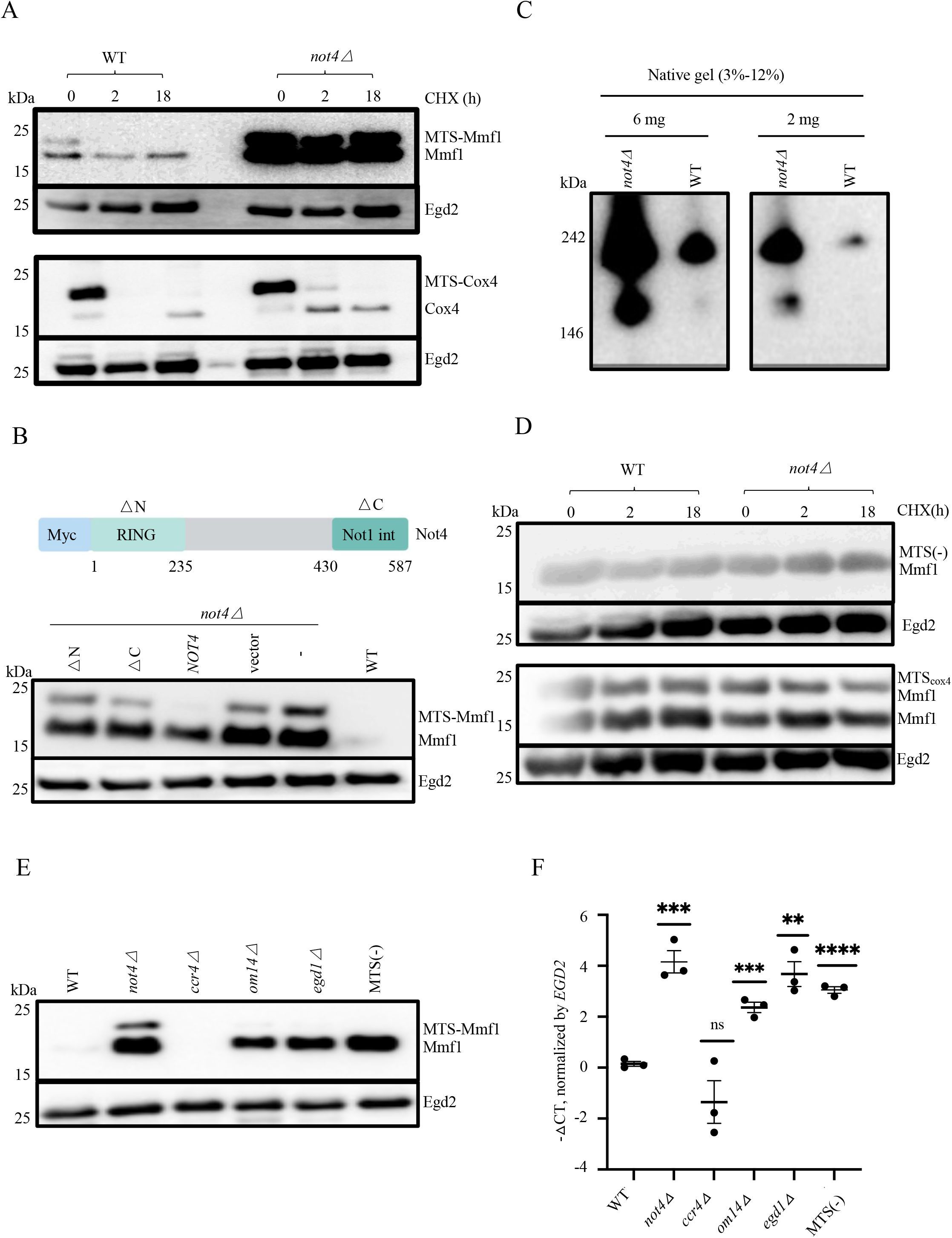
Overexpression of *MMF1* mRNA and protein in cells lacking Not4 or when Mmf1 lacks its MTS is epistatic. **A**. Analysis of the reporters was evaluated in wild type and *not4Δ* as in **Figure 1B. B**. Top: cartoon of the Myc6-Not4 coding sequence. The RING domain is located before amino acid 235 and the Not1-interaction domain is located after amino acid 430. Bottom: wild type cells (WT), *not4Δ* cells (-) or *not4Δ* cells transformed with plasmids expressing with an N-terminal Myc tag, wild type Not4, a derivative lacking the RING domain (*Δ*N) or a derivative lacking the Not1-interacting C-terminal domain (*Δ*C) and the *MMF1* reporter, were analyzed before copper induction as in **panel A**. **C**. The indicated amounts of total soluble protein extract from wild type or *not4Δ* cells expressing Mmf1 with a Taptag from its endogenous locus were analyzed by Native PAGE and western blotting with PAP antibodies. **D**. Wild type and *not4Δ* cells were analyzed for expression of the *MMF1* reporter without the MTS or with the *COX4* MTS as in **panel A**. **E and F**. Wild type and the indicated mutant cells transformed with the *MMF1* reporter or wild type cells transformed with the *MMF1* reporter without the MTS as indicated were collected at the exponential growth phase without copper induction and analyzed by western blotting with antibodies to Flag for protein levels (**E**) and by RT-qPCR for mRNA levels (**F**). The *EGD2* protein and mRNA were used as a control for loading. (**F**). The *MMF1* reporter mRNA levels were plotted to show means +/-S.E.M. of – ΔCT values. The level of significant change, relative to WT is indicated with asterisks using a non-parametric T-test (n = 3).

We tested which functional domains of Not4 were important for control of Mmf1 expression and transformed the *not4Δ* null strains carrying the reporters with plasmids encoding wild type or mutant Not4 derivatives, in particular Not4 mutants lacking their C-terminal Not1-interation domain or the N-terminal RING domain (54). Only wild type Not4 showed complementation of the Mmf1 overexpression. Notably however, the complementation from the plasmid expressing wild type Not4 was only partial (**Figure 2B**), maybe because of the presence of an N-terminal tag, or because Not4 is expressed from an episome rather than from the genomic locus.

The Not proteins are known to be important for co-translational assembly of specific protein complexes (29,30). Mmf1 forms homotrimers proposed to assemble with Mam33 trimers (40). We thus questioned whether Mmf1 complexes were appropriately formed in cells lacking Not4 and analyzed extracts of wild type and mutant cells expressing the *MMF1* reporter on native gels. Mmf1 from all strains migrated with a size between 146 and 242 kDa, larger than expected for Mmf1 homotrimers. Hence, the same apparent Mmf1 complexes could be formed in wild type cells and cells lacking Not4. However, faster migrating Mmf1 complexes were additionally seen in cells lacking Not4 (**Figure 2C**). These faster migrating Mmf1 complexes likely reflect higher expression levels of Mmf1 compared to its partner proteins, though we cannot exclude that they indicate ineffective complex assembly in mutant cells if the partner proteins are not limiting.

We next questioned whether increased expression of Mmf1 due to the absence of Not4 and the MTS were additive. However, the expression of Mmf1 without its MTS or with the Cox4 MTS was not further increased in *not4Δ* (**Figure 2D**). Hence Not4 and the Mmf1 MTS are epistatic with regard to their regulation of the Mmf1 reporter.

mRNAs that are translated with ribosome pausing can be importantly under the control of co-translational QC pathways that control both protein and mRNA levels (for review see (55)). Hence, we tested whether overexpression of the Mmf1 protein from the *MMF1* reporter in the mutants was accompanied by an overexpression of *MMF1* mRNA levels. We tested this before copper induction, when we already detect overexpression of Mmf1 protein in mutants compared to wild type (**Figure 2E**). In addition to the mutants tested above, we also tested cells lacking Ccr4, the deadenylase of the Ccr4-Not complex. The levels of the Mmf1 protein and *MMF1* mRNA were significantly higher in all mutants compared to the wild type, except in *ccr4Δ* (**Figure 2F**). Hence, expression of the *MMF1* reporter is importantly controlled co-translationally at the mRNA and protein levels, dependent upon its MTS, Not4 and the co-translational import machinery, but not upon the Ccr4 deadenylase.

### The Mmf1 MTS, co-translational import machinery and Not4 contribute to localize the *MMF1* mRNA to mitochondria

The results presented above raise the question of how Not4, the Mmf1 MTS and the co-translational import machinery together regulate expression of the *MMF1* reporter. We first questioned whether the increased Mmf1 precursor that accumulated in *not4Δ* was associated with mitochondria. By purifying mitochondria (**Figure S2A**), we detected only mature Mmf1 associated with the mitochondrial fraction (**Figure S2B**), whereas instead the precursor was detected in the cytoplasmic fraction (**Figure S2C**). From these observations we considered the possibility that the Mmf1 MTS together with Not4 might contribute to target the *MMF1* mRNA to the co-translational import machinery. To determine anchoring of the *MMF1* mRNA to the co-translational import machinery we inserted new generation MS2 stem loops (sl) (56) into the 3’UTR of the *ADH1* terminator on the reporter carrying the *MMF1* ORF, with or without its MTS (**Figure 3A**). In parallel we created a plasmid expressing the MS2-stem loop binding protein (MCP) fused C-terminally to mScarlet. We first verified that the MCP-mScarlet protein was effectively binding the mRNA. For this, we sedimented extracts from cells expressing the MCP-mScarlet with the *MMF1* reporter with or without the MS2 sl through a 60% sucrose cushion. Indeed, only in this latter case was MCP detected in the pellet from the sucrose cushion with the ribosomes (**Figure S2D**), indicating that the MCP fusion protein was effectively recruited to the *MMF1*-MS2sl mRNA.

**Figure 3.**
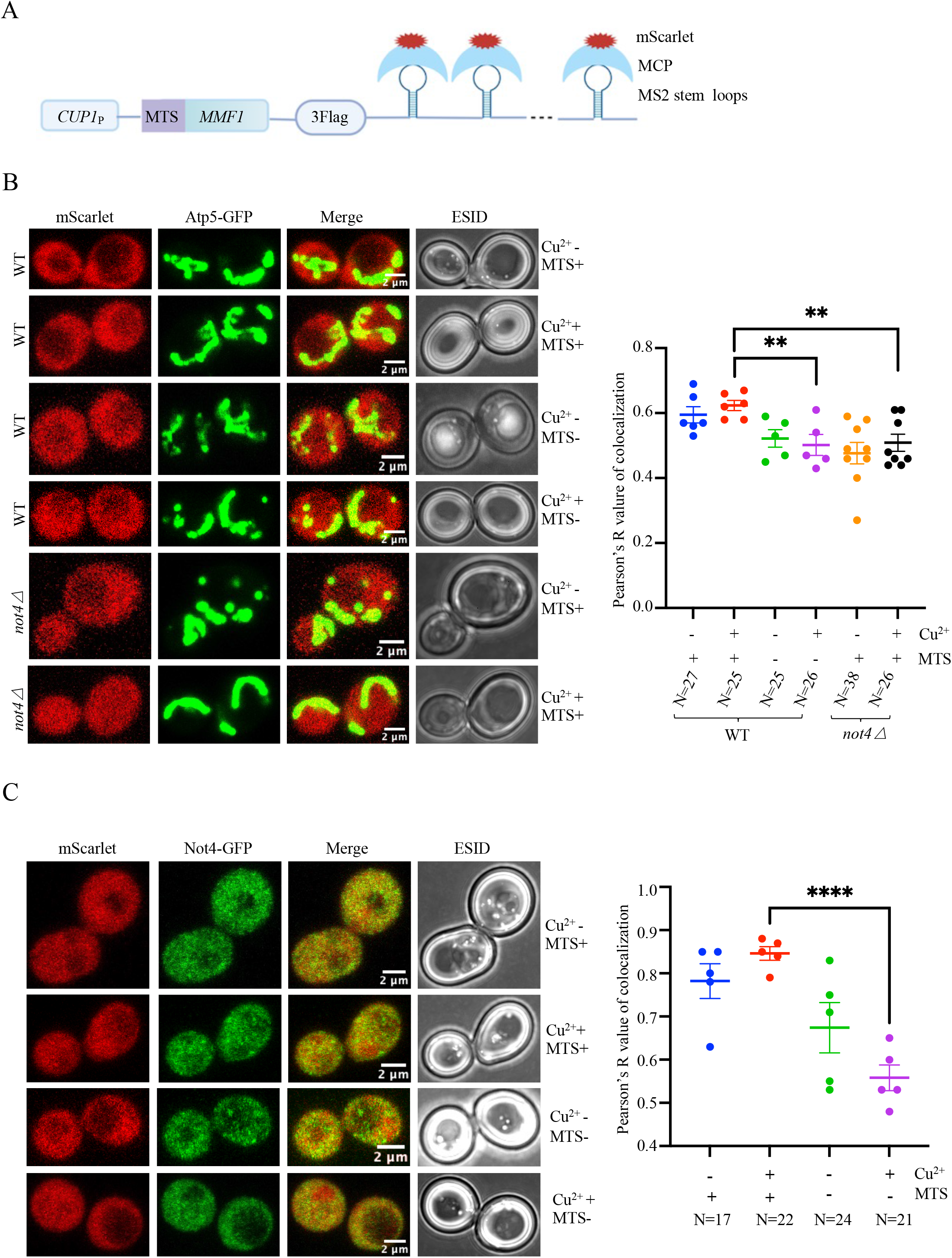
The Mmf1 MTS and Not4 contribute to the localization of the *MMF1* mRNA to the mitochondria. **A**. Cartoon of the *MMF1* reporter with inserted MS2 stem loops in the 3’UTR that can be recognized by MS2 binding protein (MCP) fused to mScarlet. **B**. Wild type cells or *not4Δ* cells as indicated expressing from its endogenous locus Atp5 fused to GFP, were transformed with the plasmid expressing MCP fused to mScarlet and the *MMF1* reporter with or without its MTS as indicated. Cells were grown to exponential phase, and induced (+) or not (-) with copper (Cu^2+^) for 10 min, then placed on agar-containing slides that were visualized at the confocal microscope to see mScarlet (left panels), Atp5-GFP (middle panels), and the merged signal (right panels), as well as the cells by phase contrast (far right panels). Representative images of 2 cells are shown. More than 25 cells were analyzed and the co-localization of the green and red signals evaluated to provide a Pearson’s R value for co-localization. **C**. Same as in panel **B** except wild type cells expressing from its own locus Not4 fused to GFP were tested.

We then transformed the plasmids in wild type or *not4Δ* strains expressing Atp5-GFP to visualize mitochondria using confocal microscopy. The MCP-mScarlet was expressed and localized all over the cytoplasm, whether the *MMF1* mRNA was induced or not (**Figure 3B**, left panels). The co-localization of the *MMF1* mRNA bound by the MCP-mScarlet (red) and mitochondrial GFP-tagged Atp5 (green) was revealed by the presence of a yellow signal, before and after copper induction (**Figure 3B**, merge panels). To evaluate the extent of this co-localization in a statistically significant way we used Prism 9 (see methods). The co-localization was similar at low or high expression of the *MMF1* reporter (before and after copper induction), and in both cases was dependent upon the MTS (**Figure 3B**, right graphic). We also analyzed colocalization of *MMF1* mRNA with Atp5-GFP before and after copper induction in cells lacking Not4. The co-localization was significantly decreased in cells lacking Not4 after copper induction (**Figure 3B**). Next, we looked at the co-localization of Not4 with the *MMF1* mRNA with or without the *MMF1* MTS, before and after copper induction, by transforming the reporter in cells expressing GFP-tagged Not4 from its endogenous locus. We noted that Not4 co-localized with the *MMF1* mRNA before and after copper induction, in a manner that was significantly dependent upon the MTS after the copper induction (**Figure 3C**).

### Egd1 ubiquitination and Caf130 limit co-translationally *MMF1* expression

The results so far indicate that the Mmf1 MTS promotes Not4 interaction with the *MMF1* mRNA. Furthermore, the MTS, Not4, as well as Egd1, regulate *MMF1* expression. Egd1 is a substrate for the Not4 ubiquitin ligase (35). We thus questioned whether ubiquitination of Egd1 by Not4 contributed to regulate *MMF1* expression. We tested expression of the *MMF1* reporter in wild type cells, or in *egd1Δ* cells transformed with either an empty vector, a vector expressing wild type Egd1, or a plasmid expressing the non-ubiquitinated Egd1_K29,30,R_ derivative (57). The Mmf1 precursor was overexpressed in *egd1Δ* as expected, and this was complemented by wild type Egd1, but not by the non-ubiquitinated Egd1 (**Figure 4A**). Notably, the addition of a plasmid expressing Egd1 to complement the absence of the genomic *EGD1* gene resulted in even less Mmf1 precursor after copper induction than when Egd1 was expressed from its endogenous locus in wild type cells.

**Figure 4.**
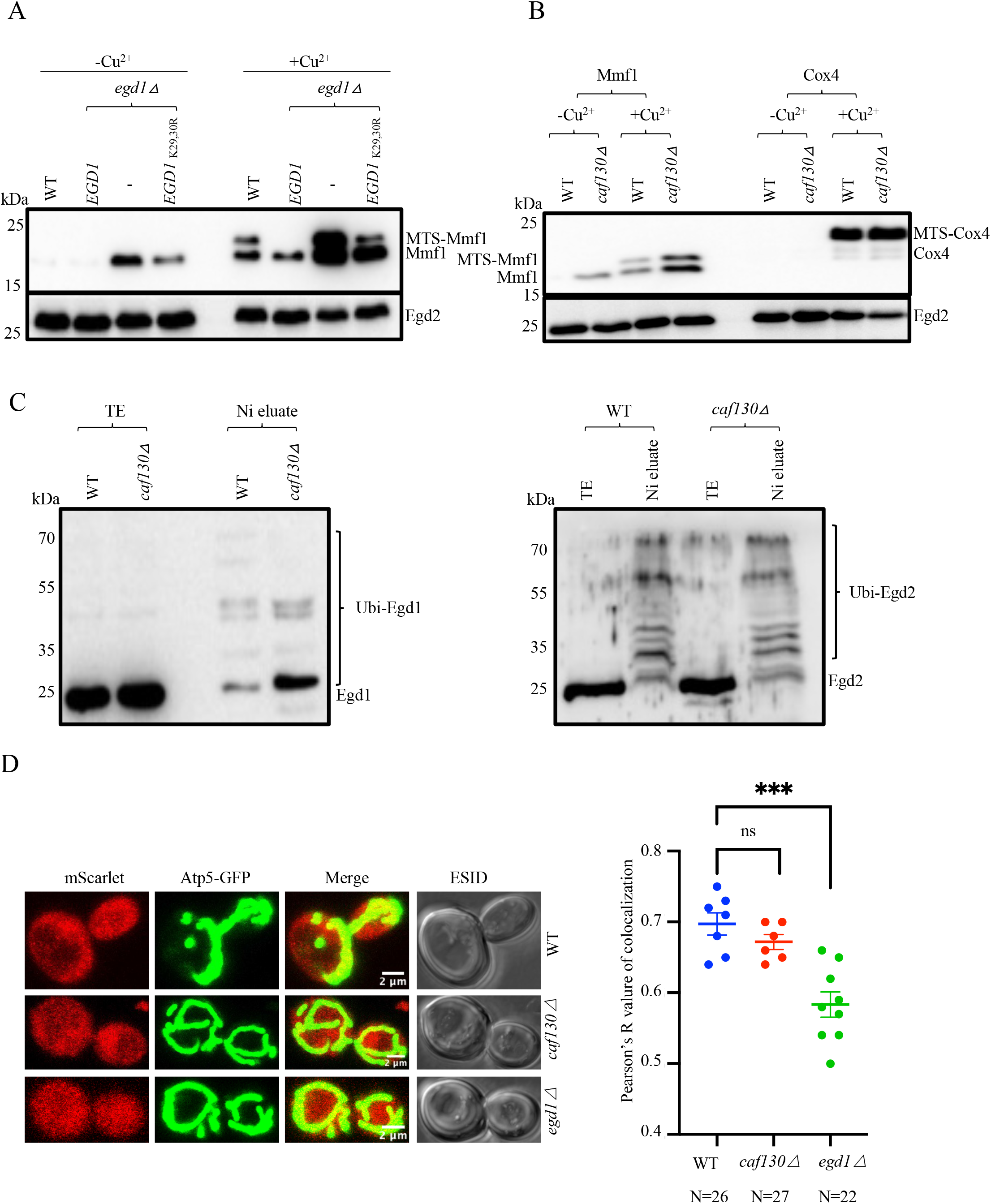
Egd1 ubiquitination and Caf130 limit expression of the *MMF1* reporter. **A**. Wild type (WT) or *egd1Δ* cells transformed with a plasmid expressing wild type HA-tagged Egd1 (Egd1), a control vector (-) or a plasmid expressing an HA-tagged Egd1 derivative that does not get ubiquitinated (Egd1_K29,30R_) were tested for expression of the *MMF1* reporter before (-Cu^2+^) and after (+Cu^2+^) a 10 min copper induction by western blotting with antibodies to Flag or Egd2 as loading control. **B**. Wild type and *caf130Δ* cells were tested for expression of the *MMF1* and *COX4* reporters as in panel **A**. **C**. Wild type and *caf130Δ* cells expressing HA-tagged Egd1 from the endogenous *EGD1* locus and transformed with a plasmid 6His-tagged ubiquitin under the *CUP1* promoter were grown in the presence of 0.1 mM CuS04. Ubiquitinated proteins were purified by nickel affinity chromatography and the presence of Egd1 in the total extract (TE) and nickel eluate (Ni-eluate) were tested for the presence of Egd1 or Egd2 with respectively antibodies to HA (left panels) or with antibodies to Egd2 (right panels) **D**. Localization of the *MMF1* mRNA was tested as in **Figure 3B** in wild type cells, *egd1Δ* and in *caf130Δ*, after a 10 min copper induction.

We have observed using Not5 affinity purification that Egd1 co-purifies with the Ccr4-Not from wild type cells, but it does not co-purify with Ccr4-Not from cells lacking Caf130, another subunit of the Ccr4-Not complex. This was also the case for the other NACß subunit, Btt1, and for the NACa subunit Egd2 (**Table S1**), as previously reported (58) and confirmed recently (59). This led us to test the expression of the Mmf1 and Cox4 reporters in cells lacking Caf130. Mmf1, but not Cox4, was overexpressed in cells lacking Caf130 (**Figure 4B**). Since Egd1 ubiquitination by Not4 was important to control Mmf1 expression, and Caf130 was important for co-purification of NAC with the Ccr4-Not complex, we determined whether ubiquitination of NAC was impaired in cells lacking Caf130. We transformed a plasmid expressing His-tagged ubiquitin from the *CUP1* promoter in *caf130Δ* cells expressing HA-tagged Egd1. After induction with copper, we affinity purified ubiquitinated proteins on a nickel resin. Total proteins and affinity-purified proteins were analyzed by western blotting for Egd1 with antibodies to HA and for Egd2 with polyclonal antibodies to Egd2 (**Figure 4C**). NAC ubiquitination was not abolished in cells lacking Caf130. However, in cells lacking Caf130 there was higher accumulation of lower molecular weight ubiquitinated forms and reduced accumulation of higher molecular weight ubiquitinated forms of Egd1, suggesting reduced turnover of ubiquitinated Egd1 in *caf130Δ*.

We then determined if Egd1 and Caf130 contributed to localize the *MMF1* mRNA to the mitochondria by confocal microscopy using the same setup described above. There was no significant change in *MMF1* mRNA co-localization with Atp5-GFP in cells lacking Caf130, whereas there was a significant decrease of this co-localization in cells lacking Egd1 (**Figure 4D**).

### RQC, as well as Cis1, Hsp104 and autophagy limit overexpression of Mmf1

As mentioned above, many QC pathways exist to avoid accumulation of proteins that arrive at the mitochondria, either overexpressed precursor proteins, mistargeted proteins or misfolded and defective proteins. Since we noted that *MMF1* but not *COX4*, was translated with ribosome pausing, we tested expression of the reporters in wild type cells and in cells lacking Hel2, a major effector of RQC. Mmf1, but not Cox4, was overexpressed in *hel2Δ*, before and after copper induction (**Figure 5A**). Mmf1 precursor and mature protein were also overexpressed in cells lacking Vms1, the tRNA hydrolase that antagonizes Rqc2 (**Figure 5B**). Cox4 expression on the other hand was not affected. We next tested the role played by components of other QC responses, starting with Cis1 that associates with the mitochondrial translocase to reduce the accumulation of mitochondrial precursor proteins. Mmf1, but not Cox4, was overexpressed in cells lacking Cis1 (**Figure 5C**), but not in cells lacking Msp1, the Cis1-interacting AAA^+^ adenosine triphosphatase (**Figure S3A**). Thus the regulation of Mmf1 overexpression involves Cis1 by mechanism distinct to “MitoCPR” (see above). Mmf1, but not Cox4, was also overexpressed in cells lacking the Hsp104 disaggregase (**Figure 5C**) or in cells lacking the exonucleases that mediate degradation of the mRNA in NGD, Ski2 or Xrn1 (60) (**Figure 5D**). We also tested whether mitophagy that removes aged and damaged mitochondria contributed to limit Mmf1 overexpression using a strain lacking Atg32, the receptor for mitophagy. However, Mmf1 levels were unaltered in *atg32 Δ* (**Figure 5E**). Mitophagy is a selective type of autophagy, so we tested whether autophagy contributed to limit Mmf1 overexpression, using cells lacking the Atg17 scaffold protein. Mmf1, but not Cox4, was overexpressed in cells lacking Atg17 (**Figure 5F**). Expression of *MMF1* without its MTS was not increased in any of the mutants of these different QC pathways (**Figure 5G**).

**Figure 5.**
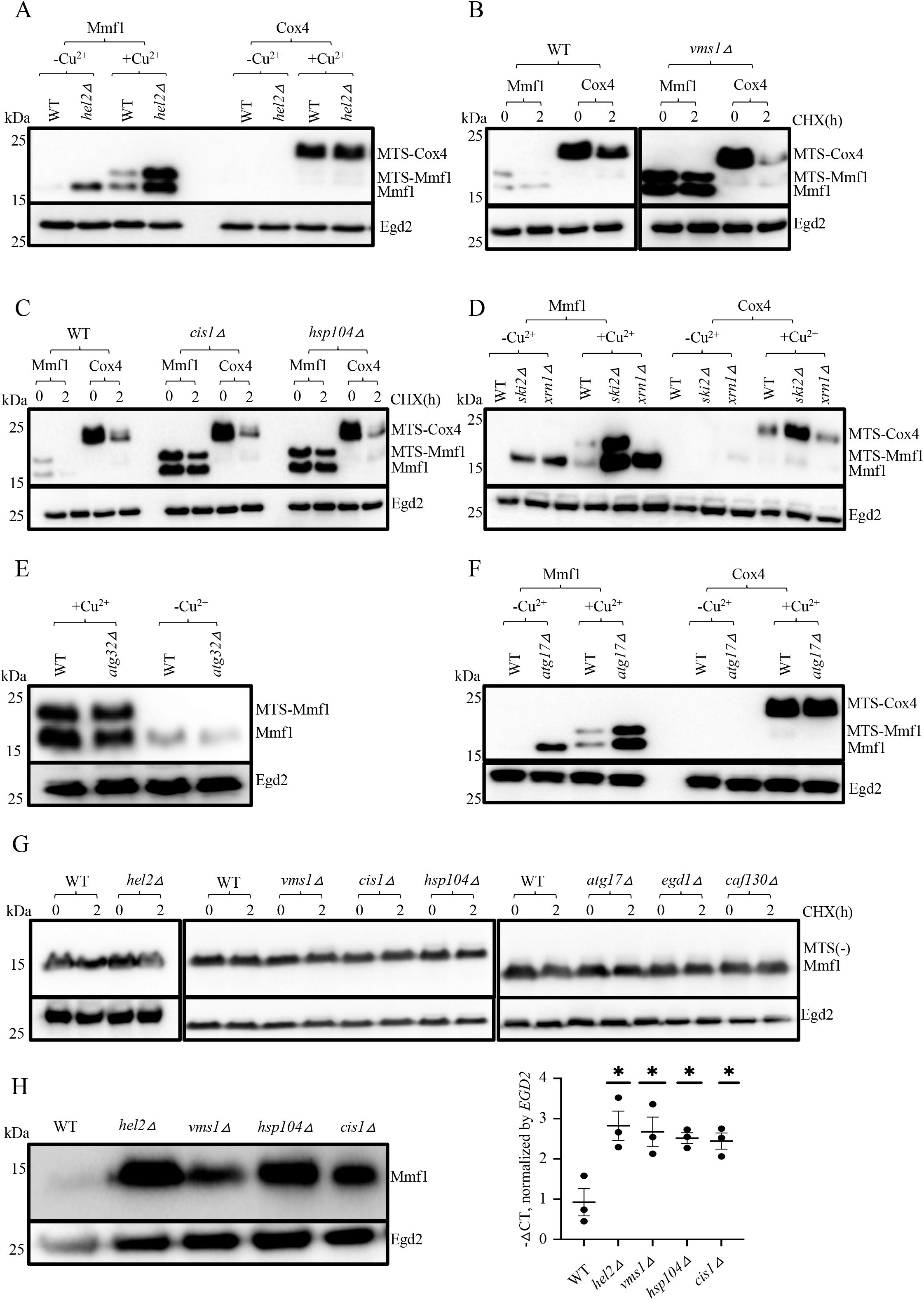
An integrated quality control response regulates expression of the *MMF1* reporter. **A**. Expression of the *MMF1* and *COX4* reporters was tested in wild type cells (WT) and in cells lacking *HEL2* as in **Figure 1B**. **B**. Expression of the*MMF1* and *COX4* reporters was tested in WT and in cells lacking *VMS1* after a 10 min copper induction (0) or 2 hours after a CHX chase (2) as in **panel A**. **C**. Same for WT and cells lacking *CIS1* or *HSP104*. **D**. Same for WT and cells lacking *SKI2* or *XRN1*. **E**. Same WT and cells lacking *ATG32*. **F**. Same for WT and cells lacking *ATG17*, before and after a 10 min copper induction. **G**. Expression of the *MMF1* reporter without MTS in WT or in cells lacking *HEL2*, *VMS1*, *CIS1*, *HSP104*, *ATG17*, *EGD1* and *CAF130* as indicated, was tested as in **panel A**. **H**. Expression of the *MMF1* reporter from cells growing exponentially without copper induction was evaluated, either the protein levels by western blotting as in **panel A** (left) or the mRNA levels (right) by RT-qPCR. For the mRNA, the levels were normalized to *EGD2* mRNA and the results are expressed as – ΔCT values in the different strains relative to WT. The level of significant change, relative to WT is indicated with asterisks using a non-parametric T-test (n = 3).

The results above indicate that ribosome pausing and many QC pathways regulate *MMF1* reporter expression, including RQC. This suggests that at least some of the response is co-translational, therefore expected to reduce protein and mRNA levels. To determine the extent of this co-translational regulation, we measured levels of *MMF1* reporter mRNA in wild type and QC mutants. The increase of Mmf1 protein in cells lacking Hel2, Vms1, Hsp104 and Cis1 (**Figure 5H**, left panel) was accompanied by an increase in the *MMF1* reporter mRNA (**Figure 5H**, right panel). In contrast, and consistent with the protein levels, the MTS-less *MMF1* reporter mRNA was not increased in *hel2Δ* (**Figure S3B**). An increase in *MMF1* mRNA also accompanied the increase of Mmf1 protein in *atg17Δ*. This was not always detectable before copper induction even when the reporter protein was increased, but very markedly and consistently detected after copper induction (**Figure S3C**). Mmf1 was increased before and after copper induction also in all other autophagy mutants tested, namely *atg11Δ, atg5Δ* and *atg8Δ* (**Figure S3D**).

These results indicate that many QC pathways work together to limit synthesis and accumulation of the Mmf1 precursor, as long as the *MMF1* ribosome nascent chain complex (RNC) is targeted to the mitochondria.

### Physiological targets of the Caf130 quality control pathway

The integrated QC mechanism determined in the experiments above is revealed with a reporter artificially overproducing the *MMF1* coding sequence in cells growing in glucose. Its relevance for endogenous *MMF1* is supported by our ribosome profiling data showing that in cells lacking Not4, endogenous *MMF1* is overexpressed and shows a defect in ribosome pausing. Besides Not4, the experiments above indicate that the Caf130 subunit of the Ccr4-Not complex is also important for this integrated QC response, and, interestingly, in cells lacking Caf130 but not in wild type cells, we detect Mam33, an interactor of Mmf1 (40) co-purifying with the Ccr4-Not complex (**Table S1**). However, *MMF1* is unlikely to be the major physiological target of this QC response in wild type cells growing in glucose, when translation of mitochondrial proteins is limited. To identify the main targets of this regulation in fermenting yeast, we performed ribosome profiling experiments (Ribo-Seq) (61) with wild type and *caf130Δ* cells growing in glucose (**Table S2**). Ribosome footprints on some mitochondrial mRNAs were up-regulated in *caf130Δ*, but *MMF1* itself was only minimally affected, and overall mitochondrial mRNAs were not significantly overrepresented in the up-regulated mRNAs (**Figure 6A**). Instead GO terms “cytoplasmic translation” “protein folding” and “metabolic processes” were the most significantly enriched GO categories within mRNAs with up-regulated ribosome footprints in *caf130Δ* (**Table S3**). Ribosomal protein mRNAs were highly significantly enriched in the upregulated group (**Figure 6B**). Interestingly, there was a very significant overlap between the mRNAs whose solubility was increased upon Not4 depletion and decreased upon Not1 depletion (see **Figure S1A**) and the mRNAs with increased ribosome footprints in *caf130Δ* (**Figure 6C**). Moreover, overall mRNAs exhibiting large increased ribosome pauses in *caf130Δ* compared to wild type cells (see methods for definition) were enriched in these overlapping mRNAs (hypergeometric test, p-value = 1.077e-06).

**Figure 6.**
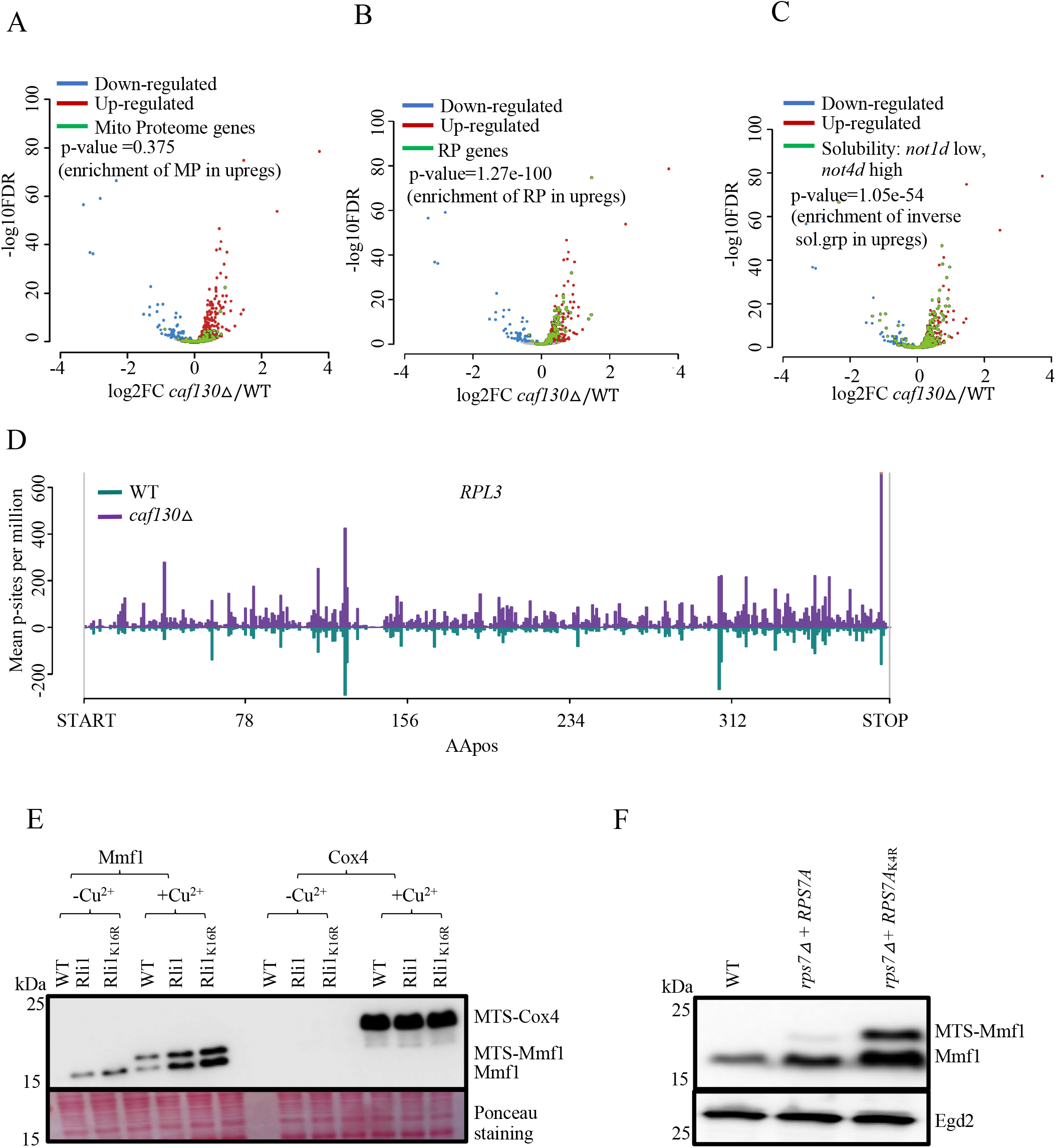
mRNAs up-regulated in *caf130Δ* are enriched for mRNAs more soluble upon Not4 depletion but less soluble upon Not1 depletion. (**A**) Volcano plot of log2FC vs -log10FDR calculated by DESeq2 comparing ribosome footprinting RPKMs in *caf130Δ* and WT. Mitochondrial proteome genes are overlaid in green and the p-value (calculated by a hypergeometric test) for their enrichment among upregulated genes is given. (**B**) as in (**A**) but with ribosomal protein gene mRNAs overlaid. (**C**) as in (**A**) but with the group of genes showing significantly changed solubility low upon Not1 depletion (*not1d*) and high upon Not4 depletion (*not4d*) as indicated in **Figure S1A** in red. (**D**) Profiles of ribosome footprints (P-site depth plots) on *RPL3* with footprints in wild type cells in green and those in *caf130Δ* in purple. The number of P-sites, per million genome wide for each sample, covering each CDS codon is calculated, averaged for each condition and plotted. **E**. Wild type cells transformed with the *MMF1* reporter were transformed with a plasmid overexpressing Rli1 or a non-complementing Rli1 derivative with 16 lysine codons mutated to arginine. The expression of the *MMF1* reporter before (-) and after (+) a 10 min copper induction (Cu^2+^) was tested by western blotting with antibodies to Flag. The ponceau staining of the blot is shown below. **F**. Expression of the *MMF1* reporter was evaluated by western blotting with antibodies to Flag, in WT and in cells and expressing wild type or non-ubiquitinated Rps7A (K4R) from a plasmid to complement the deletion of genomic *RPS7A* and *RPS7B*, grown to exponential phase and without copper induction. Antibodies to Egd2 were used to as loading control.

The *RPL3, RPL4A* and *RPL4B* mRNAs were in the group of mRNAs that showed both increased ribosome footprints in *caf130Δ* and were more soluble upon Not4 depletion, but less upon Not1 depletion. They were amongst the mRNAs with most up-regulated ribosome footprints in *caf130Δ* and showed huge increases of ribosome pausing (**Figure 6D** and **S4A**). This finding is interesting in light of a recent study indicating that Caf130 is important for a QC pathway monitoring the interaction of Rpl3 and Rpl4 nascent chains with their chaperones (59). This led us to question whether the *RPL3* and *RPL4* QC pathway might be the same as the one identified for overexpressed *MMF1*. To test this hypothesis, we compared *RPL3* and *RPL4* mRNA levels in wild type cells and in cells lacking Caf130 or Om14. The *RPL* mRNAs were overexpressed in *caf130Δ* but not in *om14Δ* (**Figure S4B**). The levels of *RPL3* and *RPL4* were also unaffected in cells lacking Hel2, Vms1, Hsp104, Cis1 and Atg17 (**Figure S4C**). This indicates that under normal conditions in glucose, *RPL3* and *RPL4* mRNAs are controlled by Caf130, but not by the other components of the QC described above. Interestingly nevertheless, the Rpl3 chaperone, Rrb1, co-purified with the Ccr4-Not complex from wild type cells but not from cells lacking Caf130 (**Table S1**). Hence, Caf130 might contribute to the delivery of the Rrb1chaperone to the Rpl3 nascent chain in the context of the Ccr4-Not complex. The dramatic increase in ribosome pausing on *RPL3* mRNA detected in *caf130Δ* (**Figure 6D**) may be the result of failed interaction of the nascent chain with the chaperone. In addition, the detection of ribosome pausing suggests that clearance of paused RNCs is not effective in *caf130Δ*.

We recently determined that Not4 ubiquitination of Rps7A and overexpression of another target of Not4 ubiquitination, Rli1, wild type or with 16 mutated lysine codons (both overexpressed Rli1 constructs are expected to lower the level of ubiquitinated Rli1 because Not4 is not co-overexpressed with Rli1), increased translation of a reporter with a stalling sequence (31). Because ribosome pausing appears relevant for the QC response that limits *MMF1* reporter, as well as *RPL3*, *RPL4A* and *RPL4B* overexpression in *caf130Δ*, we tested the impact of Rli1 overexpression before and after copper induction on expression of the *MMF1* reporter (**Figure 6E**). Mmf1 but not Cox4, was up-regulated before and after copper induction, upon Rli1 overexpression. Similarly, we tested the impact of non-ubiquitinated Rps7A on expression of the *MMF1* reporter before copper induction. Expression of Mmf1 was also increased in the non-ubiquitinated Rps7A mutant (**Figure 6F**).

## Discussion

### Targeting and pausing for quality control at the mitochondria outer membrane

In this work we show that budding yeast cells growing in glucose with limited need for mitochondria can mobilize an integrated QC response to avoid overexpression of the Mmf1 mitochondrial precursor induced from an episome. We have called this mechanism Mito-ENCay (**Figure 7**). This QC relies on the co-translational targeting of the *MMF1* mRNA to the MOM via the Mmf1 nascent chain, the Egd1 chaperone, Not4 of the Ccr4-Not complex, as well as Om14, Om45, and Tom20 (**Figure 7**, step 1). The Hsp104 disaggregase is also involved, probably to help targeting if the nascent chain starts aggregating before targeting is ensured. Then, at the MOM, ribosome pausing occurs for co-translational processes, such as folding, assembly and import, which is likely sensed by Caf130 and Cis1 (**Figure 7**, step 2). In case of defective co-translational processes, a number of factors and events (**Figure 7**, step 3), such as RQC (Vms1, Hel2), and ubiquitination of Egd1, Rps7A and Rli1 by Not4, will result in degradation of the *MMF1* mRNA to limit new Mmf1 synthesis and accumulation via both NGD (Xrn1, Ski2) and autophagy (Atg17, Atg5, Atg8 and Atg11) (**Figure 7**, step 4). Autophagy is not dependent upon the mitophagy receptor Atg32, but may be triggered by vesicles of mitochondrial fragments, with docked RNCs containing accumulated ubiquitinated factors, targets for autophagosome formation and targeting to the vacuole for degradation. The ubiquitination of RNC factors is likely to be mediated by the combined actions of Not4 and Hel2. This integrated QC pathway is not observed with MTS-less *MMF1* mRNA that is therefore overexpressed. It seems likely that mitochondrial targeting is necessary for ribosome pausing, itself necessary for the QC response. Indeed, in cells lacking Not4, targeting is defective as is ribosome pausing.

**Figure 7.**
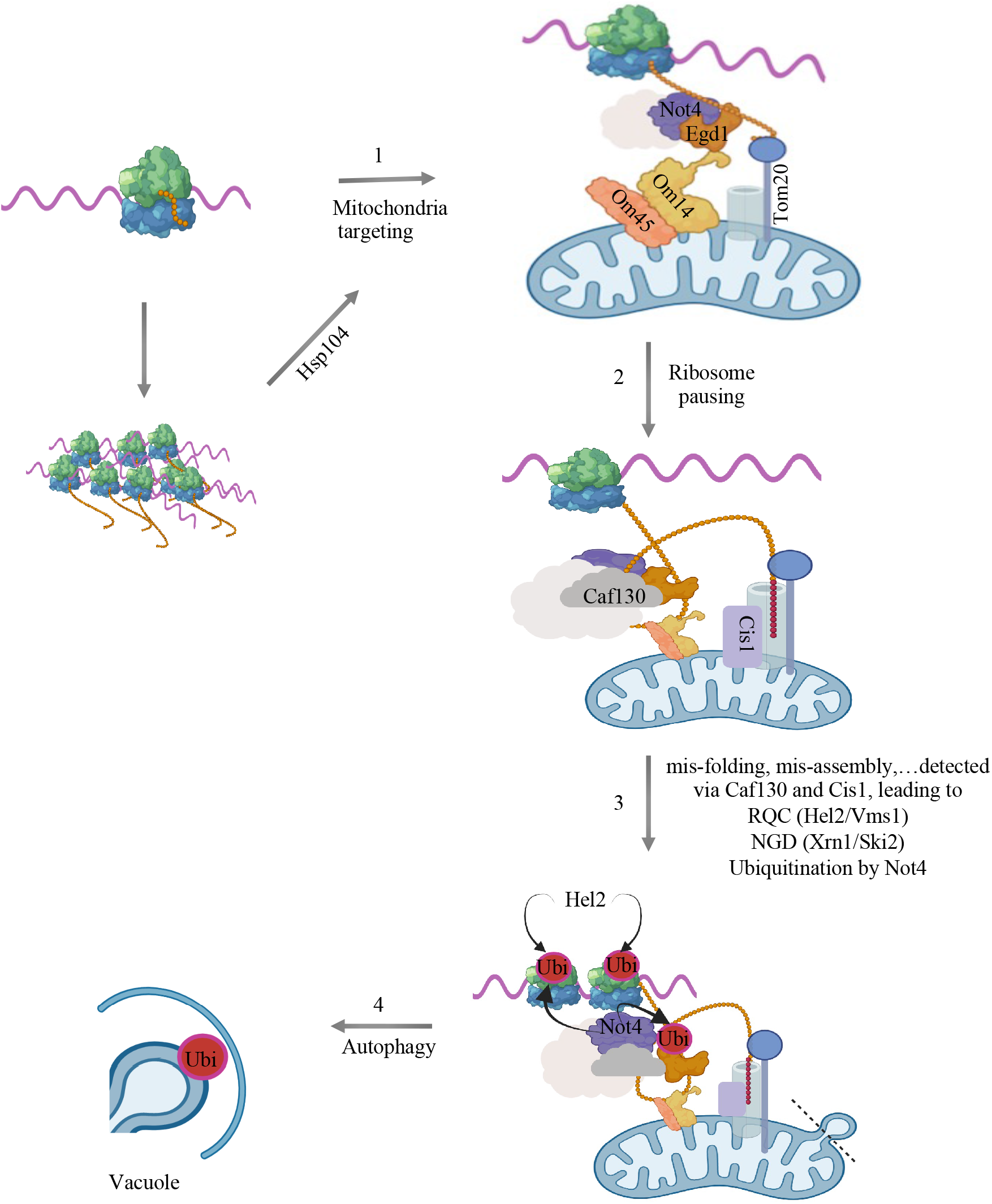
Model for limitation of translationally arrested mRNAs at the mitochondrial surface: Mito-ENCay. Overexpressed *MMF1* mRNA is targeted to the mitochondria via its nascent chain where its translation undergoes pausing, and both induction of the RQC/NGD and autophagy pathways reduce mRNA levels to limit protein synthesis and accumulation of Mmf1. This system relies on the co-translational targeting of the *MMF1* mRNA to the mitochondria via the Mmf1 nascent chain, the Egd1 chaperone, Om14, Om45 and Tom20 at the mitochondrial OM and Not4 of the Ccr4-Not complex (**step 1**). The Hsp104 disaggregase plays a regulatory role, that could be at the level of targeting, possibly if the nascent chain starts aggregating before targeting is ensured. Then, at the mitochondrial OM, ribosome pausing occurs for co-translational processes, such as folding, assembly and import, which are likely sensed by Caf130 and Cis1 (**step 2**). Caf130 may also play a scaffolding role to promote co-translational assembly. In case of defective co-translational processes, a number of factors and events (**step 3**), such as ubiquitination of Egd1, Rps7A and Rli1 by Not4, as well as RQC and ubiquitination by Hel2, will result in degradation of the *MMF1* mRNA to limit new Mmf1 synthesis and accumulation. mRNA degradation involves both NGD and autophagy, whereby vesicles of mitochondria fragments rich in OM with docked RNCs and accumulated ubiquitinated proteins, are targets for autophagosome formation and targeting to the vacuole for degradation (**step 4**).

*COX4* is not a target for Mito-ENCay. Cox4 turns over very rapidly and its production does not endanger cellular proteostasis. Therefore, Cox4 expression does not require this QC response. Nevertheless, solubility of *COX4* mRNA is also inversely regulated by Not4 and Not1, like *MMF1* mRNA, and overall ribosome footprints are increased on *COX4* mRNA in *not4Δ*. It could be that solubility of *COX4* mRNA is regulated by Not condensates (30) that might also play a role in production of Cox4, for instance for effective interaction of nascent Cox4 with cytosolic chaperones or post-translational targeting of Cox4 to mitochondria. Furthermore, *COX4* regulation by the Not proteins might depend upon 5’ or 3’UTR sequences rather than on the coding sequence as was tested in this study.

### Caf130 is at the intersection between co-translational assembly and quality control

Caf130 is a yeast specific subunit of the Ccr4-Not complex and not much is known about its function. It doesn’t contribute significantly to targeting of the *MMF1* reporter to mitochondria but from our findings seems to be part of a signaling switch from pausing to translation through pause sites. In a previous report we showed that proteasome subunits Rpt1 and Rpt2 are translated with ribosome pausing in Not condensates (that we called Not1-containing assemblysomes (NCAs)), and that when partner nascent chains interact ribosome pausing is lifted (30). Hence, Caf130 might scaffold co-translational interactions within the context of NCAs. This model is supported by our purifications of the Ccr4-Not complex from cells expressing or lacking Caf130, from which we identified many proteins co-purifying with the Ccr4-Not complex only in presence of Caf130 (see **Table S1**). These include Hsm3, a chaperone of the Rpt1 and Rpt2 proteasome assembly intermediate, and we have previously determined the role of the Ccr4-Not complex in assembly of the proteasome (30). Another example is Rba50, a chaperone involved in RNA polymerase II assembly, that we also showed previously is dependent upon the Ccr4-Not complex (27). Similarly Ada2 and Sgf29 are subunits of SAGA whose effective assembly depends upon the Ccr4-Not complex (29). Many other such factors showing Caf130-dependent co-purification with the Ccr4-Not complex are subunits of multiprotein complexes and could be targets of co-translational assembly dependent upon the Ccr4-Not complex. In addition, mRNAs with increased ribosome footprints and pausing in *caf130Δ* might be targets of this Caf130 co-translational function. For instance, Fas2 whose encoding mRNA has the third most up-regulated ribosome footprints in *caf130Δ* interacts co-translationally with Fas1 (62).

Intriguingly, several QC factors also showed Caf130-dependent co-purification with the Ccr4-Not complex. These are for instance Dcp1 and Dcp2 of the decapping complex that removes the 5’ cap structure from mRNAs prior to their degradation, Rqc2 involved in RQC, or Caf20 that competes with eIF4G for binding to eIF4E and is a repressor of translation. Caf130 might contribute to regulate translation arrest, RQC and turnover of mRNAs. This idea is compatible with our observation of accumulation of mRNAs with paused ribosomes that do not turnover in *caf130Δ*. The role of Caf130 for QC is also revealed by the fact that it is needed not for ubiquitination of Egd1 by Not4 but for turnover of ubiquitinated Egd1. An appealing idea is that prolonged or stabilized interaction of the Ccr4-Not complex at the mitochondria, mediated by interaction of Egd1 and Om14, and promoted by Caf130, is necessary for the QC. Indeed, co-localization of Not4 with the mitochondria depends upon the Mmf1 MTS whose interaction with Egd1 docks the reporter mRNA at the OM, and Caf130 mediates co-purification of Egd1 with the Ccr4-Not complex. In this context it is intriguing to mention a recent study in which it was observed that NAC subunits were amongst the most enriched proteins in polysome samples of emetine treated HCT116 cells, considered to be collided ribosomes (63). This raises the question of whether a mechanism similar to Mito-ENCay contributes to clear such collided ribosomes. Caf130 is a yeast-specific subunit of the Ccr4-Not complex, but it is possible that CNOT10 and CNOT11 interacting with the N-terminal region of CNOT1 (64) like Caf130 in yeast (59), mediate this function of Ccr4-Not in mammalian cells.

### Role of Not4 in Mito-ENCay

The Not4 subunit of the Ccr4-Not complex plays an important role in Mito-ENCay by contributing to the targeting of the Mmf1-RNC to the mitochondria, in collaboration with the nascent chain MTS and its bound chaperone Egd1. Such a function for Not4 has not previously been described, and is compatible with our previous observation that *MMF1* mRNA solubility increases upon Not4 depletion (33). In this context it is interesting to note that Not4-dependent ubiquitination of Rps7A is important for *HAC1* translational up-regulation in response to ER stress, and the presence of the *HAC1* mRNA at the ER is necessary for this up-regulation (65). *HAC1* mRNA solubility, like the solubility of *MMF1*, increases upon Not4 depletion (33). Hence, it could be that Not4 contributes to ER targeting of the *HAC1* mRNA. More globally, Not4 may generally contribute to membrane targeting of mRNAs whose solubility is increased upon Not4 depletion.

An intriguing question is how Not4 regulates mitochondrial targeting. One idea is that vesicular-mediated transport might be involved. On one hand several factors present on vesicles and related to tethering, transport or fusion of vesicles co-purify with the Ccr4-Not complex dependent upon Caf130 (such as for instance Sec2, Sec4, Sec5, Apl3 and several components of the exocyst complex (66–68)) (**Table S1**). On the other hand, components of the multi vesicular body (MVB) pathway were found to be synthetically lethal with the Not4 deletion or E3 ligase mutant (69). Recent work has revealed active mRNA transport involving transport granules that can recruit motor proteins as well as mRNAs that hitchhike on organelles, and there are close links between mRNA transport and the endocytic pathway (for reviews see (70,71)). Moreover, late endosomes have been shown to be sites of local translation important for mitochondrial maintenance in axons (72). It was recently shown that condensates of Tis11, an RNA binding protein that can associate with the Ccr4-Not complex, is important for translation in proximity to the ER (73). This supports a possible role of NCAs in mRNA targeting to vesicles and/or organelles, that would explain the regulation of mRNA solubility by Not1 and Not4 that we have recently identified (33), including *MMF1* studied here.

Not4 plays a role in the QC response beyond mRNA localization, via its ubiquitination of Egd1, Rps7A and Rli1. Hel2 ubiquitinates ribosomal proteins in response to collided ribosomes (34,74), including Rps7A first mono-ubiquitinated by Not4. It could be that Hel2 can similarly polyubiquitinate Egd1 and Rli1 after Not4 mono-ubiquitination in specific QC conditions. Protein ubiquitination is necessary in many types of selective autophagy as a mark for cargo recognition and a signal for process initiation by recruitment of specific autophagy adaptor proteins (also known as autophagy receptors) (for review see (75)). In addition, a recent study proposed a role for Not4’s ubiquitination of Rli1 in the context of paused RNCs at the MOM for mitophagy in flies (76). Thus, it seems likely that in yeast multiple ubiquitination events by Not4 and Hel2 can mark the RNCs at the MOM for autophagy as a backup when NGD is overwhelmed.

## Accession of data

All data has been deposited in public data bases. The Ribo-Seq GEO for WT and *not4Δ* is GSE137613, RNA-Seq GEO for WT, *not1d, not4d* is GSE168290 and the GEO accession for *caf130D* ribosome profiling is GSE206973. The mass spectrometry proteomics data have been deposited to the ProteomeXchange Consortium via the PRIDE (77) partner repository with the dataset identifier PXD035672 and 10.6019/PXD035672.

## Acknowledgments

We would like to thank Agnieszka Chacinska for providing us with the backbone of the reporter plasmid and the laboratories of Mafalda Escobar Henriques, Wolf Dieter, Lars Steimetz, Rachel Green, Marc Hochstrasser, Koji Okamoto and Shuh-ichi Nishikawa for many strains. We thank Denis Martinvalet and Ravish Rashpa for preliminary observations, ideas and technical support, Laetitia Maillard and Szabolcs Zahoran for experimental support. We thank the Bioimaging platform of the Faculty of Medicine of the University of Geneva for technical help with microscopy and image processing and the BioRender software that was used to create the figures.

---

**Figure S1.**
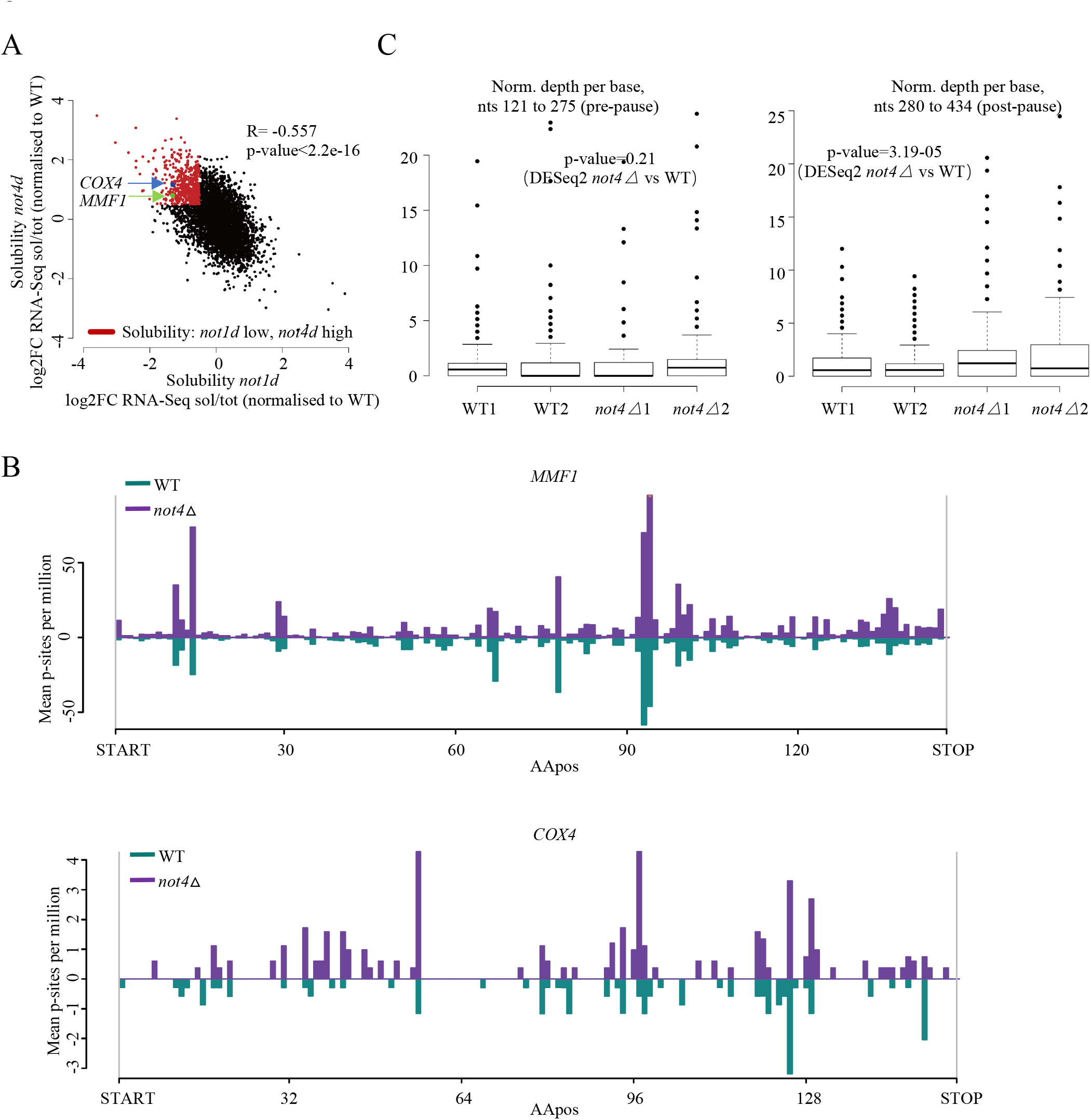
*MMF1* and *COX4* mRNAs are more soluble upon Not4 depletion and less upon Not1 depletion. **A**. Indication of *MMF1* and *COX4* mRNAs in a scatterplot from (doi:https://doi.org/10.1101/2022.03.14.484207) comparing changes in mRNA solubilities, corrected to WT, before and after Not1 and Not4 depletion. mRNAs significantly more soluble upon Not4 depletion and less upon Not1 depletion are labelled in red. **B**. Profiles of ribosome footprints (P-site depth plots) on *MMF1* (upper panel) and *COX4* (lower panel) with footprints in wild type cells in green and those in *not4Δ* in purple. The number of P-sites, per million genome wide for each sample, covering each CDS codon with corresponding amino acid position indicated (AApos) is calculated, averaged for each condition and plotted. **C**. Quantification of mRNA footprints in wild type and *not4Δ* duplicate samples on equal segments of the mRNA before (left) and after (right) the apparent ribosome pausing site. Boxplots of P-sites per million for each base of the *MMF1* CDS in WT and *not4Δ* for the region between the large pause and the stop codon (nucleotides 280-434, right panel) and an equally-sized region just upstream of the pause (nucleotides 121-275). Only the region post-pause shows significant changes in counts (DESeq2 p-value = 3.19e-5).

**Figure S2.**
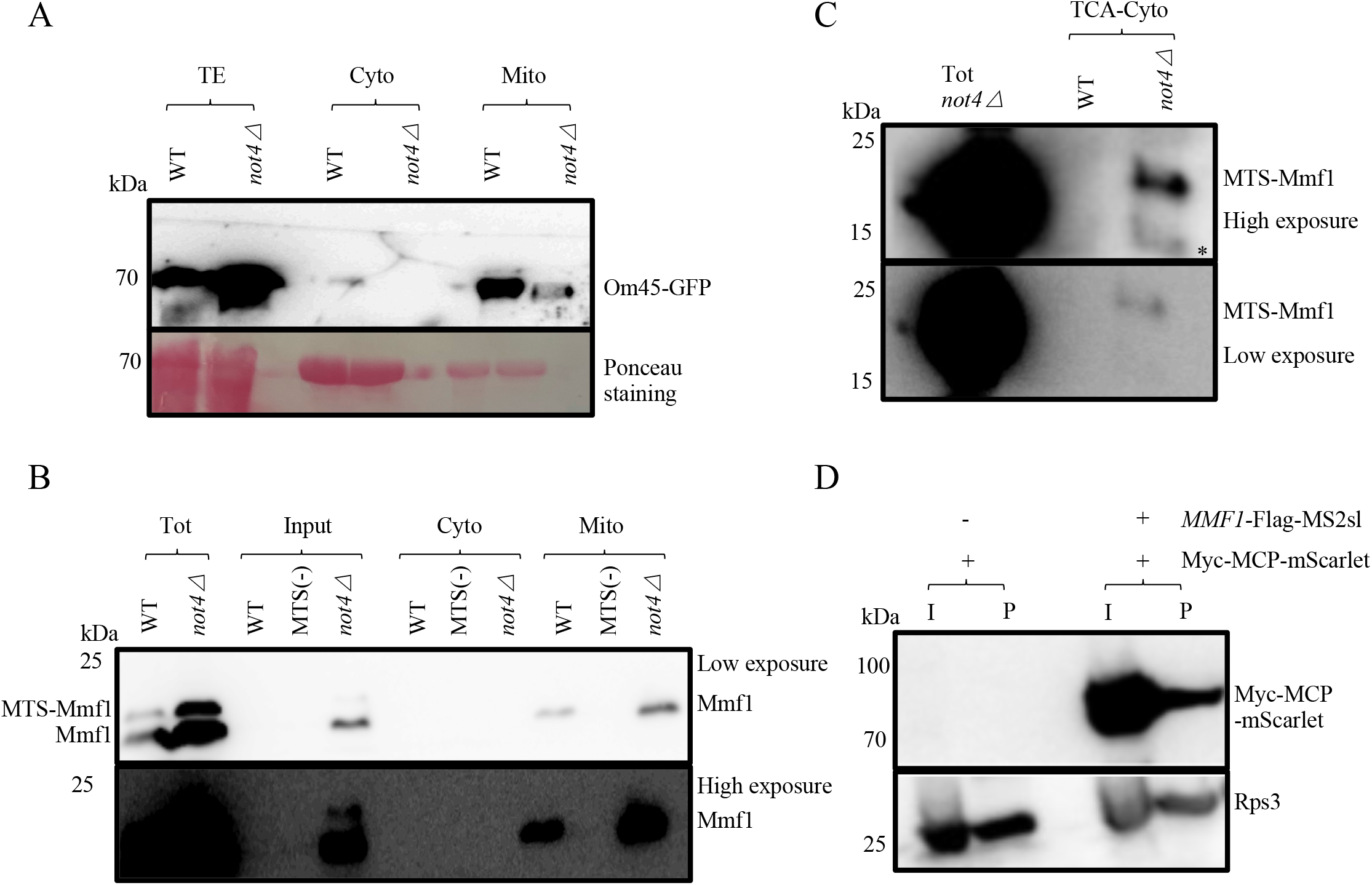
The overexpressed Mmf1 precursor accumulates in the cytoplasm in *not4Δ*. **A**. Extracts from wild type and *not4Δ* cells expressing Om45-GFP from its own locus were prepared and fractionated using a commercial kit. The total extract (TE), then cytosolic (Cyto) and mitochondrial (Mito) fractions were analyzed by western blotting with a GFP antibody, to verify the fractionation approach. **B**. The same experiment as in panel **A** was performed with WT or *not4Δ* cells overexpressing the *MMF1* reporter with or without an MTS sequence (MTS-) as indicated, after a 10 min copper induction. The western blot was revealed with antibodies to Flag and a low (upper panel) and high (lower panel) exposure is shown. **C**. The total extract from *not4Δ* and the cytosolic fractions from WT and *not4Δ* shown in panel **B**, were TCA precipitated for concentration and resuspended for analysis by western blotting with antibodies to Flag. A high (upper panel) and low (lower panel) exposure are shown. An unrelated signal is indicated by *. **D**. Cells expressing Myc-MCP-mScarlet with or without the *MMF1* reporter with MS2 stem loops (MS2sl) were lysed and the total extract (I) was sedimented on a 60% sucrose cushion. The ribosome-containing pellet (P) was analyzed by western blotting for the presence of the Myc-MCP-mScarlet fusion by western blotting with antibodies to Myc. The levels of Rps3 were analyzed as a control.

**Figure S3.**
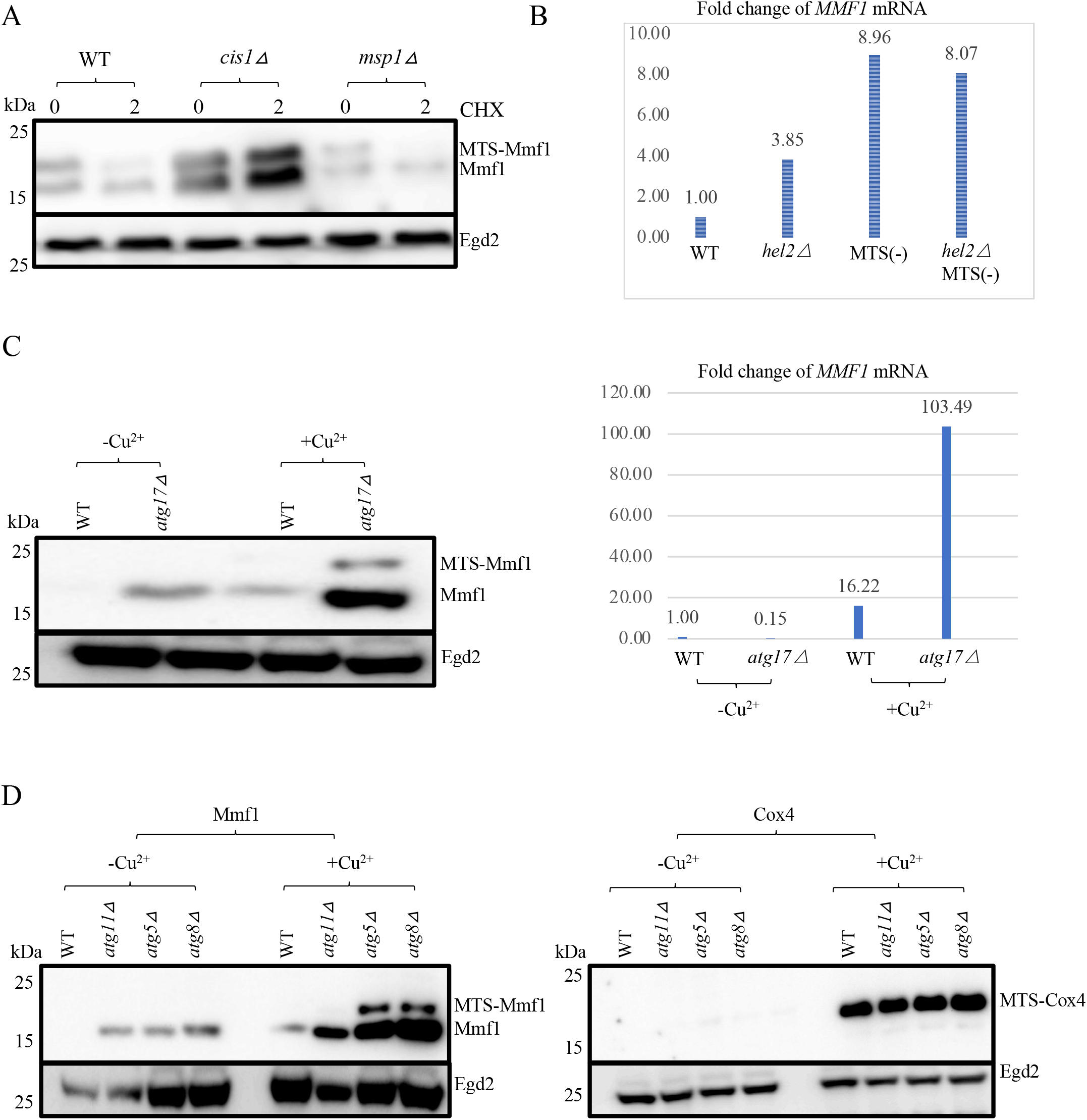
*MMF1* reporter is upregulated in autophagy mutants but not in cells lacking Msp1. **A**. Expression of the *MMF1* reporter was tested in WT, *cis1Δ* and *msp1Δ* as in **Figure 5C**. **B**. Levels of the *MMF1* and MTS-less *MMF1* reporter mRNA were evaluated in wild type cells or cells lacking Hel2 as indicated. The reporter levels were normalized to *EGD2* mRNA and the results are expressed as fold change in the different strains relative to WT. One significant experiment is shown. **C**. Expression of the *MMF1* reporter was tested in WT and *atg17Δ* at both the protein (left) and mRNA (right) level, before (-Cu^2+^) and after (+Cu^2+^) copper induction. One significant experiment is shown. **D**. Mmf1 (left panel) and Cox4 (right panel) levels were tested in WT, *atg11Δ, atg5Δ* and *atg8Δ*, before (-Cu^2+^) and after (+Cu^2+^) copper induction as in **Figure 5D**.

**Figure S4.**
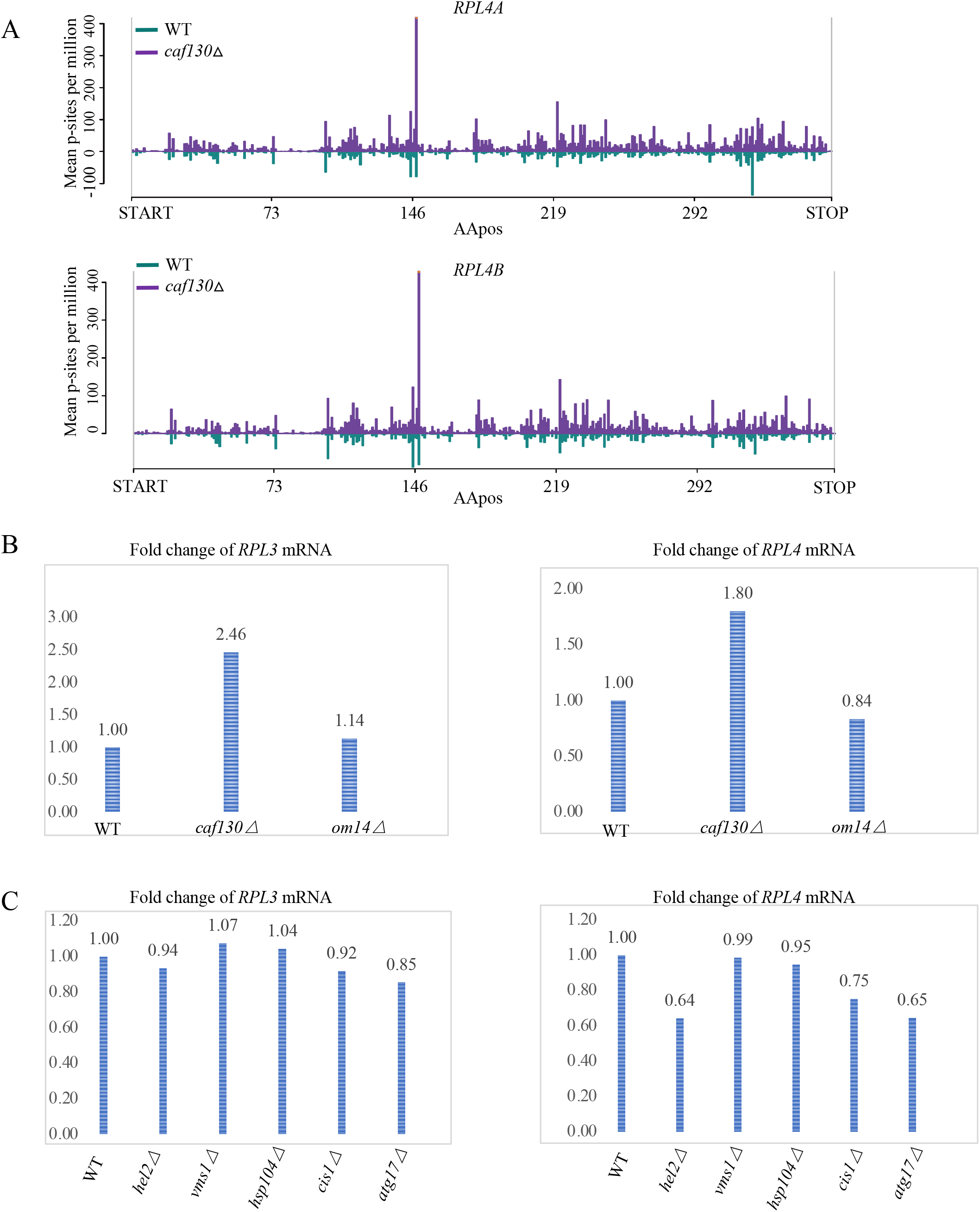
*RPL4* mRNA levels and ribosome pausing are up-regulated in *caf130Δ*. **A**. Profiles of ribosome footprints (P-site depth plots) on *RPL4A* and *RPL4B* with footprints in wild type cells in green and those in *caf130Δ* in purple. The number of P-sites, per million genome wide for each sample, covering each CDS codon is calculated, averaged for each condition and plotted. **B, C**. *RPL3* and *RPL4* mRNA levels were evaluated in (**B**) WT, *caf130Δ* and *om14Δ* and (**C**) WT, *hel2Δ, vms1Δ, hsp104Δ, cis1Δ* and *atg17Δ*, by RT-qPCR. *EGD2* was the loading control. The mRNA levels were normalized to *EGD2* mRNA and the results are expressed as fold change in the different strains relative to WT. One significant experiment is shown.

## Supplementary table legends

**Table S1. Some proteins co-purify with the Ccr4-Not complex only in presence, others only in absence, of Caf130**. Tandem affinity chromatography was performed from total protein extracts from wild type or *caf130Δ* cells expressing Tap-tagged Not5, Not1 or Caf1 proteins expressed from their endogenous locus, as indicated. The purified proteins were identified by LC-MS/MS and a score for the presence of proteins is provided (see materials and methods). Several proteins were detected only in purifications from wild type cells (in all, top panel, in 3 out of 4, second panel and 2 out of 4 in the third panel). In the fourth panel, 2 proteins were detected only in *caf130Δ*. For Mam33, it was detected only in the purifications from *caf130Δ* by 2 peptides for one of the purifications and by 1 peptide for the 2 others. Proteins mentioned in the manuscript are highlighted.

**Table S2. Ribosome footprinting from wild type and *caf130Δ***

**Table S3. GO-term analysis of mRNAs with increased ribosome footprints in *caf130Δ***

**Table S4. Strains, plasmids, oligonucleotides and antibodies used in this work**

## References

1. Chandel, N.S. (2014) Mitochondria as signaling organelles. BMC Biol, 12, 34.

2. Friedman, J.R. and Nunnari, J. (2014) Mitochondrial form and function. Nature, 505, 335–343.

3. Nunnari, J. and Suomalainen, A. (2012) Mitochondria: in sickness and in health. Cell, 148, 1145–1159.

4. Farrar, G.J., Chadderton, N., Kenna, P.F. and Millington-Ward, S. (2013) Mitochondrial disorders: aetiologies, models systems, and candidate therapies. Trends Genet, 29, 488–497.

5. Williams, C.C., Jan, C.H. and Weissman, J.S. (2014) Targeting and plasticity of mitochondrial proteins revealed by proximity-specific ribosome profiling. Science, 346, 748–751.

6. Saint-Georges, Y., Garcia, M., Delaveau, T., Jourdren, L., Le Crom, S., Lemoine, S., Tanty, V., Devaux, F. and Jacq, C. (2008) Yeast mitochondrial biogenesis: a role for the PUF RNA-binding protein Puf3p in mRNA localization. PLoS One, 3, e2293.

7. Gerber, A.P., Herschlag, D. and Brown, P.O. (2004) Extensive association of functionally and cytotopically related mRNAs with Puf family RNA-binding proteins in yeast. PLoS Biol, 2, E79.

8. Lesnik, C., Cohen, Y., Atir-Lande, A., Schuldiner, M. and Arava, Y. (2014) OM14 is a mitochondrial receptor for cytosolic ribosomes that supports co-translational import into mitochondria. Nat Commun, 5, 5711.

9. Lesnik, C., Golani-Armon, A. and Arava, Y. (2015) Localized translation near the mitochondrial outer membrane: An update. RNA Biol, 12, 801–809.

10. Ahmed, A.U. and Fisher, P.R. (2009) Import of nuclear-encoded mitochondrial proteins: a cotranslational perspective. Int Rev Cell Mol Biol, 273, 49–68.

11. Kershaw, C.J., Costello, J.L., Talavera, D., Rowe, W., Castelli, L.M., Sims, P.F., Grant, C.M., Ashe, M.P., Hubbard, S.J. and Pavitt, G.D. (2015) Integrated multi-omics analyses reveal the pleiotropic nature of the control of gene expression by Puf3p. Sci Rep, 5, 15518.

12. Burri, L., Vascotto, K., Gentle, I.E., Chan, N.C., Beilharz, T., Stapleton, D.I., Ramage, L. and Lithgow, T. (2006) Integral membrane proteins in the mitochondrial outer membrane of Saccharomyces cerevisiae. FEBSJ, 273, 1507–1515.

13. Reimann, B., Bradsher, J., Franke, J., Hartmann, E., Wiedmann, M., Prehn, S. and Wiedmann, B. (1999) Initial characterization of the nascent polypeptide-associated complex in yeast. Yeast, 15, 397–407.

14. Rospert, S., Dubaquie, Y. and Gautschi, M. (2002) Nascent-polypeptide-associated complex. Cell Mol Life Sci, 59, 1632–1639.

15. del Alamo, M., Hogan, D.J., Pechmann, S., Albanese, V., Brown, P.O. and Frydman, J. (2011) Defining the specificity of cotranslationally acting chaperones by systematic analysis of mRNAs associated with ribosome-nascent chain complexes. PLoS Biol, 9, e1001100.

16. George, R., Walsh, P., Beddoe, T. and Lithgow, T. (2002) The nascent polypeptide-associated complex (NAC) promotes interaction of ribosomes with the mitochondrial surface in vivo. FEBS Lett, 516, 213–216.

17. Wiedemann, N. and Pfanner, N. (2017) Mitochondrial Machineries for Protein Import and Assembly. Annu Rev Biochem, 86, 685–714.

18. Avendano-Monsalve, M.C., Ponce-Rojas, J.C. and Funes, S. (2020) From cytosol to mitochondria: the beginning of a protein journey. Biol Chem, 401, 645–661.

19. Zurita Rendon, O., Fredrickson, E.K., Howard, C.J., Van Vranken, J., Fogarty, S., Tolley, N.D., Kalia, R., Osuna, B.A., Shen, P.S., Hill, C.P. et al. (2018) Vms1p is a release factor for the ribosome-associated quality control complex. Nat Commun, 9, 2197.

20. Izawa, T., Park, S.H., Zhao, L., Hartl, F.U. and Neupert, W. (2017) Cytosolic Protein Vms1 Links Ribosome Quality Control to Mitochondrial and Cellular Homeostasis. Cell, 171, 890–903 e818.

21. Onishi, M., Yamano, K., Sato, M., Matsuda, N. and Okamoto, K. (2021) Molecular mechanisms and physiological functions of mitophagy. EMBO J, 40, e104705.

22. Miller, J.E. and Reese, J.C. (2012) Ccr4-Not complex: the control freak of eukaryotic cells. Crit Rev Biochem Mol Biol, 47, 315–333.

23. Denis, C.L. and Chen, J. (2003) The CCR4-NOT complex plays diverse roles in mRNA metabolism. Prog Nucleic Acid Res Mol Biol, 73, 221–250.

24. Collart, M.A. (2003) Global control of gene expression in yeast by the Ccr4-Not complex. Gene, 313, 1–16.

25. Collart, M.A. and Panasenko, O.O. (2012) The Ccr4-Not complex. Gene, 492, 42–53.

26. Collart, M.A. (2016) The Ccr4-Not complex is a key regulator of eukaryotic gene expression. Wiley Interdiscip Rev RNA, 7, 438–454.

27. Villanyi, Z., Ribaud, V., Kassem, S., Panasenko, O.O., Pahi, Z., Gupta, I., Steinmetz, L., Boros, I. and Collart, M.A. (2014) The Not5 subunit of the ccr4-not complex connects transcription and translation. PLoS Genet, 10, e1004569.

28. Gupta, I., Villanyi, Z., Kassem, S., Hughes, C., Panasenko, O.O., Steinmetz, L.M. and Collart, M.A. (2016) Translational Capacity of a Cell Is Determined during Transcription Elongation via the Ccr4-Not Complex. Cell Rep, 15, 1782–1794.

29. Kassem, S., Villanyi, Z. and Collart, M.A. (2017) Not5-dependent co-translational assembly of Ada2 and Spt20 is essential for functional integrity of SAGA. Nucleic Acids Res, 45, 1186–1199.

30. Panasenko, O.O., Somasekharan, S.P., Villanyi, Z., Zagatti, M., Bezrukov, F., Rashpa, R., Cornut, J., Iqbal, J., Longis, M., Carl, S.H. et al. (2019) Co-translational assembly of proteasome subunits in NOT1-containing assemblysomes. Nat Struct Mol Biol, 26, 110–120.

31. Allen, G.E., Panasenko, O.O., Villanyi, Z., Zagatti, M., Weiss, B., Pagliazzo, L., Huch, S., Polte, C., Zahoran, S., Hughes, C.S. et al. (2021) Not4 and Not5 modulate translation elongation by Rps7A ubiquitination, Rli1 moonlighting, and condensates that exclude eIF5A. Cell Rep, 36, 109633.

32. Buschauer, R., Matsuo, Y., Sugiyama, T., Chen, Y.H., Alhusaini, N., Sweet, T., Ikeuchi, K., Cheng, J., Matsuki, Y., Nobuta, R. et al. (2020) The Ccr4-Not complex monitors the translating ribosome for codon optimality. Science, 368.

33. Allen, G., Weiss, B., Panasenko, O., Huch, S., Villanyi, Z., Albert, B., Dilg, D., Zagatti, M., Schaughency, P., Liao, S.E. et al. (2022) Not1 and Not4 inversely determine mRNA solubility that sets the dynamics of co-translational events. bioRxiv.

34. Ikeuchi, K., Tesina, P., Matsuo, Y., Sugiyama, T., Cheng, J., Saeki, Y., Tanaka, K., Becker, T., Beckmann, R. and Inada, T. (2019) Collided ribosomes form a unique structural interface to induce Hel2-driven quality control pathways. EMBO J, 38.

35. Panasenko, O., Landrieux, E., Feuermann, M., Finka, A., Paquet, N. and Collart, M.A. (2006) The yeast Ccr4-Not complex controls ubiquitination of the nascent-associated polypeptide (NAC-EGD) complex. J Biol Chem, 281, 31389–31398.

36. Goldstrohm, A.C., Hook, B.A., Seay, D.J. and Wickens, M. (2006) PUF proteins bind Pop2p to regulate messenger RNAs. Nat Struct Mol Biol, 13, 533–539.

37. Goldstrohm, A.C., Seay, D.J., Hook, B.A. and Wickens, M. (2007) PUF protein-mediated deadenylation is catalyzed by Ccr4p. J Biol Chem, 282, 109–114.

38. Jackson, J.S., Jr., Houshmandi, S.S., Lopez Leban, F. and Olivas, W.M. (2004) Recruitment of the Puf3 protein to its mRNA target for regulation of mRNA decay in yeast. RNA, 10, 1625–1636.

39. Quenault, T., Lithgow, T. and Traven, A. (2011) PUF proteins: repression, activation and mRNA localization. Trends Cell Biol, 21, 104–112.

40. Pu, Y.G., Jiang, Y.L., Ye, X.D., Ma, X.X., Guo, P.C., Lian, F.M., Teng, Y.B., Chen, Y. and Zhou, C.Z. (2011) Crystal structures and putative interface of Saccharomyces cerevisiae mitochondrial matrix proteins Mmf1 and Mam33. J Struct Biol, 175, 469–474.

41. Roloff, G.A. and Henry, M.F. (2015) Mam33 promotes cytochrome c oxidase subunit I translation in Saccharomyces cerevisiae mitochondria. Mol Biol Cell, 26, 2885–2894.

42. Longtine, M.S., McKenzie, A., Demarini, D.J., Shah, N.G., Wach, A., Brachat, A., Philippsen, P. and Pringle, J.R. (1998) Additional modules for versatile and economical PCR-based gene deletion and modification in *Saccharomyces cerevisiae*. Yeast, 14, 953–961.

43. Wittig, I., Braun, H.P. and Schagger, H. (2006) Blue native PAGE. Nat Protoc, 1, 418–428.

44. Azzouz, N., Panasenko, O.O., Colau, G. and Collart, M.A. (2009) The CCR4-NOT complex physically and functionally interacts with TRAMP and the nuclear exosome. PLoS One, 4, e6760.

45. Keller, A., Nesvizhskii, A.I., Kolker, E. and Aebersold, R. (2002) Empirical statistical model to estimate the accuracy of peptide identifications made by MS/MS and database search. Anal Chem, 74, 5383–5392.

46. Nesvizhskii, A.I., Keller, A., Kolker, E. and Aebersold, R. (2003) A statistical model for identifying proteins by tandem mass spectrometry. Anal Chem, 75, 4646–4658.

47. Pfaffl, M.W. (2001) A new mathematical model for relative quantification in real-time RT-PCR. Nucleic Acids Res, 29, e45.

48. Martin, M. (2011) Cutadapt Removes Adapter Sequences from High-Throughput Sequencing Reads. EMBnet Journal, 17, 10–12.

49. Kim, D., Langmead, B. and Salzberg, S.L. (2015) HISAT: a fast spliced aligner with low memory requirements. Nat Methods, 12, 357–360.

50. Langmead, B. and Salzberg, S.L. (2012) Fast gapped-read alignment with Bowtie 2. Nat Methods, 9, 357–359.

51. Lauria, F., Tebaldi, T., Bernabo, P., Groen, E.J.N., Gillingwater, T.H. and Viero, G. (2018) riboWaltz: Optimization of ribosome P-site positioning in ribosome profiling data. PLoS Comput Biol, 14, e1006169.

52. Love, M.I., Huber, W. and Anders, S. (2014) Moderated estimation of fold change and dispersion for RNA-seq data with DESeq2. Genome Biol, 15, 550.

53. Fan, A.C. and Young, J.C. (2011) Function of cytosolic chaperones in Tom70-mediated mitochondrial import. Protein Pept Lett, 18, 122–131.

54. Panasenko, O.O. and Collart, M.A. (2012) Presence of Not5 and ubiquitinated Rps7A in polysome fractions depends upon the Not4 E3 ligase. Molecular microbiology, 83, 640–653.

55. Ikeuchi, K., Izawa, T. and Inada, T. (2018) Recent Progress on the Molecular Mechanism of Quality Controls Induced by Ribosome Stalling. Front Genet, 9, 743.

56. Pichon, X., Robert, M.C., Bertrand, E., Singer, R.H. and Tutucci, E. (2020) New Generations of MS2 Variants and MCP Fusions to Detect Single mRNAs in Living Eukaryotic Cells. Methods Mol Biol, 2166, 121–144.

57. Panasenko, O.O., David, F.P. and Collart, M.A. (2009) Ribosome association and stability of the nascent polypeptide-associated complex is dependent upon its own ubiquitination. Genetics, 181, 447–460.

58. Cui, Y., Ramnarain, D.B., Chiang, Y.C., Ding, L.H., McMahon, J.S. and Denis, C.L. (2008) Genome wide expression analysis of the CCR4-NOT complex indicates that it consists of three modules with the NOT module controlling SAGA-responsive genes. Mol Genet Genomics, 279, 323–337.

59. Pillet, B., Mendez-Godoy, A., Murat, G., Favre, S., Stumpe, M., Falquet, L. and Kressler, D. (2022) Dedicated chaperones coordinate co-translational regulation of ribosomal protein production with ribosome assembly to preserve proteostasis. Elife, 11.

60. Inada, T. (2013) Quality control systems for aberrant mRNAs induced by aberrant translation elongation and termination. Biochim Biophys Acta, 1829, 634–642.

61. Ingolia, N.T., Ghaemmaghami, S., Newman, J.R. and Weissman, J.S. (2009) Genome-wide analysis in vivo of translation with nucleotide resolution using ribosome profiling. Science, 324, 218–223.

62. Shiber, A., Doring, K., Friedrich, U., Klann, K., Merker, D., Zedan, M., Tippmann, F., Kramer, G. and Bukau, B. (2018) Cotranslational assembly of protein complexes in eukaryotes revealed by ribosome profiling. Nature, 561, 268–272.

63. Sinha, N.K., Ordureau, A., Best, K., Saba, J.A., Zinshteyn, B., Sundaramoorthy, E., Fulzele, A., Garshott, D.M., Denk, T., Thoms, M. et al. (2020) EDF1 coordinates cellular responses to ribosome collisions. Elife, 9.

64. Mauxion, F., Preve, B. and Seraphin, B. (2013) C2ORF29/CNOT11 and CNOT10 form a new module of the CCR4-NOT complex. RNA Biol, 10, 267–276.

65. Matsuki, Y., Matsuo, Y., Nakano, Y., Iwasaki, S., Yoko, H., Udagawa, T., Li, S., Saeki, Y., Yoshihisa, T., Tanaka, K. et al. (2020) Ribosomal protein S7 ubiquitination during ER stress in yeast is associated with selective mRNA translation and stress outcome. Sci Rep, 10, 19669.

66. Wiederkehr, A., De Craene, J.O., Ferro-Novick, S. and Novick, P. (2004) Functional specialization within a vesicle tethering complex: bypass of a subset of exocyst deletion mutants by Sec1p or Sec4p. J Cell Biol, 167, 875–887.

67. Guo, W., Roth, D., Walch-Solimena, C. and Novick, P. (1999) The exocyst is an effector for Sec4p, targeting secretory vesicles to sites of exocytosis. EMBO J, 18, 1071–1080.

68. Yeung, B.G., Phan, H.L. and Payne, G.S. (1999) Adaptor complex-independent clathrin function in yeast. Mol Biol Cell, 10, 3643–3659.

69. Mulder, K.W., Inagaki, A., Cameroni, E., Mousson, F., Winkler, G.S., De Virgilio, C., Collart, M.A. and Timmers, H.T. (2007) Modulation of Ubc4p/Ubc5p-mediated stress responses by the RING-finger-dependent ubiquitin-protein ligase Not4p in Saccharomyces cerevisiae. Genetics, 176, 181–192.

70. Muntjes, K., Devan, S.K., Reichert, A.S. and Feldbrugge, M. (2021) Linking transport and translation of mRNAs with endosomes and mitochondria. EMBO Rep, e52445.

71. Das, S., Vera, M., Gandin, V., Singer, R.H. and Tutucci, E. (2021) Intracellular mRNA transport and localized translation. Nat Rev Mol Cell Biol, 22, 483–504.

72. Cioni, J.M., Lin, J.Q., Holtermann, A.V., Koppers, M., Jakobs, M.A.H., Azizi, A., Turner-Bridger, B., Shigeoka, T., Franze, K., Harris, W.A. et al. (2019) Late Endosomes Act as mRNA Translation Platforms and Sustain Mitochondria in Axons. Cell, 176, 56–72 e15.

73. Ma, W. and Mayr, C. (2018) A Membraneless Organelle Associated with the Endoplasmic Reticulum Enables 3’UTR-Mediated Protein-Protein Interactions. Cell, 175, 1492–1506 e1419.

74. Juszkiewicz, S., Chandrasekaran, V., Lin, Z., Kraatz, S., Ramakrishnan, V. and Hegde, R.S. (2018) ZNF598 Is a Quality Control Sensor of Collided Ribosomes. Mol Cell, 72, 469–481 e467.

75. Chen, R.H., Chen, Y.H. and Huang, T.Y. (2019) Ubiquitin-mediated regulation of autophagy. J Biomed Sci, 26, 80.

76. Wu, Z., Wang, Y., Lim, J., Liu, B., Li, Y., Vartak, R., Stankiewicz, T., Montgomery, S. and Lu, B. (2018) Ubiquitination of ABCE1 by NOT4 in Response to Mitochondrial Damage Links Co-translational Quality Control to PINK1-Directed Mitophagy. Cell Metab, 28, 130–144 e137.

77. Perez-Riverol, Y., Bai, J., Bandla, C., Garcia-Seisdedos, D., Hewapathirana, S., Kamatchinathan, S., Kundu, D.J., Prakash, A., Frericks-Zipper, A., Eisenacher, M. et al. (2022) The PRIDE database resources in 2022: a hub for mass spectrometry-based proteomics evidences. Nucleic Acids Res, 50, D543–D552.

## References in Table S4 (1–6)

1. Kassem, S., Villanyi, Z. and Collart, M.A. (2017) Not5-dependent co-translational assembly of Ada2 and Spt20 is essential for functional integrity of SAGA. Nucleic Acids Res, 45, 1186–1199.

2. Panasenko, O.O. and Collart, M.A. (2012) Presence of Not5 and ubiquitinated Rps7A in polysome fractions depends upon the Not4 E3 ligase. Molecular microbiology, 83, 640–653.

3. Panasenko, O.O., David, F.P. and Collart, M.A. (2009) Ribosome association and stability of the nascent polypeptide-associated complex is dependent upon its own ubiquitination. Genetics, 181, 447–460.

4. Panasenko, O.O., Somasekharan, S.P., Villanyi, Z., Zagatti, M., Bezrukov, F., Rashpa, R., Cornut, J., Iqbal, J., Longis, M., Carl, S.H. et al. (2019) Co-translational assembly of proteasome subunits in NOT1-containing assemblysomes. Nat Struct Mol Biol, 26, 110–120.

5. Hope, I. and Struhl, K. (1986) Functional dissection of a eukaryotic transcriptional activator protein. Cell, 46, 885–894.

6. Azzouz, N., Panasenko, O.O., Colau, G. and Collart, M.A. (2009) The CCR4-NOT complex physically and functionally interacts with TRAMP and the nuclear exosome. PLoS One, 4, e6760.

